# Cytochrome oxidase requirements in *Bordetella* reveal insights into evolution towards life in the mammalian respiratory tract

**DOI:** 10.1101/2024.02.29.582880

**Authors:** Liliana S. McKay, Alexa R. Spandrio, Richard M. Johnson, M. Ashley Sobran, Sara A. Marlatt, Katlyn B. Mote, Margaret R. Dedloff, Zachary M. Nash, Steven M. Julio, Peggy A. Cotter

**Author notes:** CONTRIBUTIONS Conceptualization: Liliana S. McKay, M. Ashley Sobran, Peggy A. Cotter Data curation: Liliana S. McKay, M. Ashley Sobran Formal analysis: Liliana S. McKay, M. Ashley Sobran, Peggy A. Cotter Funding acquisition: Steven M. Julio, Peggy A. Cotter Investigation: Liliana S. McKay, Alexa R. Spandrio, Richard M. Johnson, M. Ashley Sobran, Sara A. Marlatt, Katlyn B. Mote, Margaret R. Dedloff, Zachary M. Nash, Steven M. Julio, Peggy A. Cotter Methodology: Liliana S. McKay Software: N/A Supervision: Steven M. Julio, Peggy A. Cotter Validation: Liliana S. McKay, Peggy A. Cotter Visualization: Liliana S. McKay, Peggy A. Cotter Writing – original draft: Liliana S. McKay, Peggy A. Cotter Writing – review & editing: Liliana S. McKay, Peggy A. Cotter, Richard M. Johnson, Alexa R. Spandrio.

## Abstract

Little is known about oxygen utilization during infection by bacterial respiratory pathogens. The classical *Bordetella* species, including *B. pertussis*, the causal agent of human whooping cough, and *B. bronchiseptica*, which infects nearly all mammals, are obligate aerobes that use only oxygen as the terminal electron acceptor for electron transport-coupled oxidative phosphorylation. *B. bronchiseptica*, which occupies many niches, has eight distinct cytochrome oxidase-encoding loci, while *B. pertussis*, which evolved from a *B. bronchiseptica*-like ancestor but now only survives only in and between human respiratory tracts, has only three functional cytochrome oxidase-encoding loci: *cydAB1, ctaCDFGE1*, and *cyoABCD1*. To test the hypothesis that the three cytochrome oxidases encoded within the *B. pertussis* genome represent the minimum number and class of cytochrome oxidase required for respiratory infection, we compared *B. bronchiseptica* strains lacking one or more of the eight possible cytochrome oxidases *in vitro* and *in vivo*. No individual cytochrome oxidase was required for growth in ambient air, and all three of the cytochrome oxidases conserved in *B. pertussis* were sufficient for growth in ambient air and low oxygen. Using a high-dose, large-volume persistence model and a low-dose, small-volume establishment of infection model, we found that *B. bronchiseptica* producing only the three *B. pertussis*-conserved cytochrome oxidases was indistinguishable from the wild-type strain for infection. We also showed that CyoABCD1 is sufficient to cause the same level of bacterial burden in mice as the wild-type strain and is thus the primary cytochrome oxidase required for murine infection, and that CydAB1 and CtaCDFGE1 fulfill auxiliary roles or are important for aspects of infection we have not assessed, such as transmission. Our results shed light on respiration requirements for bacteria that colonize the respiratory tract, the environment at the surface of the ciliated epithelium, and the evolution of virulence in bacterial pathogens.

**AUTHOR SUMMARY:** Cytochrome oxidases, critical components for aerobic respiration, have been shown to be vital for pathogenesis and tissue tropism in several bacterial species. However, the majority of the research has focused on facultative anaerobes and infections of microoxic to anaerobic host environments, like the gut. We sought to understand the role of cytochrome oxidases during respiratory infection by *Bordetella bronchiseptica*, an obligate aerobe, performing the first analysis of cytochrome oxidases in an extracellular respiratory pathogen that we know of. By comparing *B. bronchiseptica* to the closely related *B. pertussis*, a strictly human-specific pathogen and the causative agent of whooping cough, we found three cytochrome oxidases that are important for growth and survival within the mammalian respiratory tract. We also found that a *bo_3_*-type cytochrome oxidase, predicted to have a low affinity for oxygen and therefore best suited to ambient air levels of oxygen, was sufficient for both the establishment of infection and persistence in the respiratory tract in mice. Our findings reveal the importance of low affinity cytochrome oxidases in respiratory pathogens, and emphasize the need to study the physiology of diverse pathogens.

## INTRODUCTION

To be effective pathogens, bacteria must be able to survive within the host. Even pathogens that survive exclusively within a host must contend with diverse microenvironments, moving from the initial site of infection to colonization sites and eventually to a new host while avoiding or neutralizing host defenses. As bacteria encounter new microenvironments within the host, they must adjust their physiologies to continue to grow. Most of the research on bacterial pathogens has focused on identifying and characterizing virulence factors. However, there is now an increased appreciation for the role that metabolism and physiology play during infection. Without energy, bacteria cannot survive and therefore cannot cause disease.

Electron transport-coupled oxidative phosphorylation pairs the transfer of electrons from donors to acceptors with the transport of protons across a membrane, generating the proton motive force used to drive the synthesis of ATP from ADP. This process occurs in a step-wise manner via the electron transport chain (ETC), starting with electron donors like NADPH and ending with a terminal electron acceptor. Aerobic respiration uses oxygen as the terminal electron acceptor, but alternative terminal electron acceptors like nitrate or nitrite are used by some bacteria to grow anaerobically. Redox-active protein complexes with heme cofactors called cytochrome oxidases are responsible for the transfer of electrons to oxygen. Cytochrome oxidases are named based on the types of hemes (*a, b, c, d,* or *o*) that the proteins bind. *bo_3_*-type, *aa_3_*-type, and *cbb_3_*-type cytochrome oxidases are all heme-copper oxidases (HCOs), so-named for the copper ions that form a bi-metallic center with hemes in their catalytic subunits, while *bd*-type cytochrome oxidases form a unique class of heme-only cytochrome oxidases found exclusively in bacteria and archaea with no sequence homology to HCOs [1,2]. Cytochrome oxidases can also be categorized by their electron donors. Quinol oxidases receive their electrons directly from reduced quinones, while cytochrome *c* oxidases receive their electrons from cytochrome *c*. Cytochrome oxidases can also be grouped according to their affinity for oxygen: either low-affinity (K_m_ for O_2_ ≅ 200nM) [3], which are best suited for atmospheric levels of oxygen (∼21%), or high-affinity, which can function in microoxic conditions (K_m_ for O_2_ = 3-8 nM) [4,5]. Increased affinity for oxygen comes with a trade-off; high-affinity cytochrome oxidases generate less proton motive force per electron transferred than low-affinity cytochrome oxidases [6].

Respiration as a whole, and cytochrome oxidases in particular, have been shown to be required for infection in many pathogenic species, including gastrointestinal pathogens *Escherichia coli* [7,8]*, Vibrio cholerae* [9]*, Listeria monocytogenes* [10,11], *Shigella flexneri* [12,13], and *Salmonella enterica* Serovar Typhimurium [14–16]; the oral pathogen *Aggregatibacter actinomycetemcomitans* [17]; the respiratory pathogen *Mycobacterium tuberculosis* [18,19]; and pathogens that can cause many types of infection including bacteremia and sepsis like *Staphylococcus aureus* [20] and group B *Streptococcus* [21,22]. However, the majority of pathogens assessed have been facultative anaerobes, which do not require cytochrome oxidases for survival. Additionally, many of these pathogens colonize microenvironments that are expected to have limited levels of oxygen.

The classical *Bordetella* species (*B. pertussis, B. parapertussis_Hu_,* and *B. bronchisceptica*) are extremely closely related bacteria that colonize the upper and lower respiratory tracts of humans (in the case of *B. pertussis* and *B. parapertussis_Hu_*) and other mammals (in the case of *B. bronchiseptica*). They are obligate aerobes that use only oxygen as their terminal electron acceptor for electron transport-coupled oxidative phosphorylation [23,24]. Therefore, they depend solely on components of the ETC, like cytochrome oxidases, to generate energy. Previous work in *B. pertussis* has shown that cytochrome *c* is not required for survival *in vitro* [25]. Little is known, however, about the role of cytochrome oxidases in the classical *Bordetella* species.

In this study, we leveraged the evolution of classical bordetellae to investigate which cytochrome oxidases are important for respiration in the mammalian respiratory tract, using *B. bronchiseptica* as our model organism. *B. bronchiseptica*, which occupies multiple environmental niches, has the potential to produce eight distinct cytochrome oxidases. By contrast, *B. pertussis*, which survives in the environment only during transmission between hosts, has undergone significant genome reduction as it evolved from a *B. bronchiseptica*-like ancestor [24]. Genomic comparisons between *B. bronchiseptica* and *B. pertussis* found that *B. pertussis* strains have only broadly conserved three intact cytochrome oxidase-encoding gene loci. We hypothesized that the three cytochrome oxidases conserved in *B. pertussis* represent the minimum number and class of cytochrome oxidases required for bordetellae to survive within, and briefly between, hosts. We genetically manipulated our bacteria to determine the necessity and sufficiency of different cytochrome oxidases, first for growth *in vitro* under ambient air and different atmospheric conditions, then during infection, using two different mouse models of respiratory infection. Our results demonstrate that *B. bronchiseptica* is able to survive with as few as one cytochrome oxidase, even during infection and when challenged with low oxygen conditions. These findings also inform our understanding of the murine respiratory tract, indicating that bacteria may have more access to oxygen while attached to the ciliated epithelium than previously supposed.

## RESULTS

### *Bordetella bronchiseptica* has eight cytochrome oxidase-encoding gene loci

The *B. bronchiseptica* RB50 genome has eight cytochrome oxidase-encoding gene loci, annotated computationally based on predicted amino acid homology (**Table 1**). Three of the loci are predicted to encode high-affinity cytochrome oxidases. *ccoNOQP* (abbreviated *cco*) encodes a *cbb_3_-type* cytochrome oxidase. In *cbb_3_*-type cytochrome oxidases, CcoN contains the active site that facilitates the reduction of molecular oxygen to water and thus functions as the catalytic subunit [26,27]. Two loci encode *bd*-type cytochrome oxidases (*cydAB1* and *cydAB3*, abbreviated *cyd1* and *cyd3*). The *cyd1* operon of *B. bronchiseptica* also includes a third subunit, *cydX*, which is seen in some, but not all, *bd-*type cytochrome oxidases and may be important for complex function. The variation across different species in number of subunits, structural arrangement of coordinated hemes, and positioning of the quinol binding site makes it difficult to draw definitive conclusions about *bd-*type cytochrome oxidases [28]. CydA is the subunit that coordinates the three hemes required by this cytochrome oxidase, making it the catalytic subunit; however, *bd-type* cytochrome oxidases require both CydA and CydB for catalytic activity, as evidenced by the loss of *d-*type heme when the genes encoding either subunit are deleted [29,30].

**Table 1.** Cytochrome oxidase-encoding gene loci in classical *Bordetella* species.

| Class | Gene names | *Bb* locus (RB50) | Predicted oxygen affinity | Is the locus present/intact?<sup>1</sup> |  |  |  |
| --- | --- | --- | --- | --- | --- | --- | --- |
|  |  |  |  | Complex I (n=6) | Complex II (n=14) | Complex III (n=4) | Complex IV (n=4)<sup>3</sup> |
| *cbb<sub>3</sub>* | *ccoNOQP* | BB3329-3326 | High | 100% | 0% | 100% | 75% |
| *bd* | *cydAB1* | BB4498-4497 | High | 100% | 100% | 100% | 100% |
|  | *cydAB3* | BB4012-4011 | High | 100% | 50% | 100% | 100% |
|  | *cioAB* | BB1238-1239 | Low<sup>2</sup> | 100% | 0% | 100% | 75% |
| *aa<sub>3</sub>* | *ctaCDFGE1* | BB4831-4827 | Low | 100% | 100% | 100% | 100% |
|  | *ctaD2* | BB4674 | Low | 100% | 7.1% | 100% | 100% |
| *bo<sub>3</sub>* | *cyoABCD1* | BB1283-1286 | Low | 100% | 92.9% | 100% | 75% |
|  | *cyoABCD2* | BB1310-1307 | Low | 100% | 0% | 100% | 75% |

Shaded cells represent cytochrome oxidase-encoding loci that are broadly conserved in *B. pertussis* (Complex II)
<sup>1</sup> Further information about strains compared can be found in Supplemental Table 1; details about the differences from RB50 can be found in Supplemental Table 2.
<sup>2</sup> *cioAB*, formerly known as *cydAB2*, was previously predicted to encode a high-affinity cytochrome oxidase.
<sup>3</sup> One strain, the environmental isolate HT200, has gained mutations in 4 cytochrome oxidase-encoding gene loci; the other three strains have maintained all 8 loci.

Five of the eight cytochrome oxidase-encoding loci are predicted to encode low-affinity cytochrome oxidases. These include one complete *aa_3_*-type cytochrome oxidase (*ctaCDFGE1,* abbreviated *cta1*) and one lone *aa_3_*-type subunit (*ctaD2*). The *cta1* operon contains two additional genes: a putative membrane protein (BB4829, here named *ctaF1*) and a putative DUF2970-containing membrane protein (BB4828, here named *ctaG1*). This operon structure is also found in *Achromobacter* species, which are closely related to *Bordetella*. *ctaF1* shares no homology to the gene of the same name found in *Bacillus* species, which encodes part of a *caa3*-type cytochrome oxidase [31]. CtaD is the catalytic subunit in *aa_3_*-type cytochrome oxidases [32]. While CtaD2 cannot function within the electron transport chain on its own, it could be forming heterocomplexes with CtaC1 and CtaE1, as CcoN orphan subunits have been shown to do in *Pseudomonas aeruginosa* [33]. Two loci encode *bo_3_*-type cytochrome oxidases (*cyoABCD1* and *cyoABCD2*, abbreviated *cyo1* and *cyo2*). CyoB is the catalytic subunit for *bo_3_*-type cytochrome oxidases [34,35]. Finally, the *cioAB* locus (abbreviated *cio*) encodes a cyanide-insensitive oxidase (CIO). CIOs are a subclass of *bd-*type cytochrome oxidases that are low-affinity and were first characterized in *P. aeruginosa,* where CioAB confers resistance to cyanide [36]. CIOs require both subunits to be functional, like other *bd*-type cytochrome oxidases.

### *Bordetella pertussis* has only three broadly conserved cytochrome oxidase-encoding gene loci

Genomic comparisons of the classical bordetellae have found that the three subspecies can be divided into four complexes [37]. Complex I, which includes RB50, is most similar to the last common ancestor shared by the classical bordetellae and contains *B. bronchiseptica* strains, primarily those isolated from non-human mammals. Complex II contains *B. pertussis* and Complex III contains *B. parapertussis_Hu_*. Complex IV, which phylogenetically lies between Complex I and II, contains *B. bronchiseptica* strains, primarily those isolated from humans. A recent genotyping study identified a novel *B. bronchiseptica* branch, named lineage II, that is distinct from the classical *Bordetella* [38]. This branch primarily contains strains that were formerly classed as Complex IV and had been identified by other groups as being divergent from other classical bordetellae [39,40].

*B. pertussis* and *B. parapertussis_Hu_* have shrunken their genomes since evolving from a *B. bronchiseptica*-like ancestor, with genomes approximately 77% and 89% the size of the *B. bronchiseptica* genome, respectively [24]. Additionally, *B. pertussis* and *B. parapertussis_Hu_* have specialized, infecting only humans instead of maintaining the wide mammalian host range and environmental survival capabilities of *B. bronchiseptica*. We investigated which cytochrome oxidase-encoding genes have been conserved across the classical bordetellae, hypothesizing that the human-adapted *B. pertussis* and *B. parapertussis_Hu_* have lost cytochrome oxidases that are not required for survival within, and briefly between, human respiratory tracts. We chose 28 *Bordetella* strains for comparison, including both laboratory strains and clinical isolates: 6 Complex I, 14 Complex II, 4 Complex III, and 4 Complex IV (2 of which could be classified as lineage II) **(S1 Table).** We found that almost all the strains in Complex I, III, and IV have maintained intact copies of all eight cytochrome oxidase-encoding gene loci (**Table 1**). Only one strain, the environmental isolate HT200, differed, having acquired frameshift mutations leading to premature stop codons in *ccoN*, *cioB*, *cyoB1,* and *cyoB2* which most likely would prevent the complexes encoded by each of these cytochrome oxidase-encoded loci (here notated as Cco, Cio, Cyo1, and Cyo2) from functioning in this strain **(S2 Table).** This result is intriguing, as it indicates that not all cytochrome oxidases are required to survive in the environment, or at least within the thermal spring in which HT200 was isolated [40]. This strain is also a lineage II strain, and thus phylogenetically distant from the classical *Bordetella* strains [38] **(S1 Figure)**.

*B. pertussis* strains have significantly diverged from the other classical bordetellae in terms of their cytochrome oxidase-encoding genes. All 14 examined strains contained premature stop codons within *ccoN* and *cyoA2* **(S2 Table**). These mutations would most likely prevent Cco and Cyo2 from functioning. Interestingly, the point mutations that led to the premature stops were identical across all 14 strains, indicating that these mutations were acquired by the ancestral strain that became *B. pertussis.* Additionally, all 14 strains have lost *cio* and 13 strains have lost *ctaD2*. The identical changes across diverse *B. pertussis* strains indicate that functional copies of Cco, Cio, Cyo2, and CtaD2 were lost during the evolution from the last common ancestor shared by *B. pertussis* and *B. bronchiseptica*.

There are mutations in cytochrome oxidase-encoding genes that differ between *B. pertussis* strains. Strain 18323, a common laboratory strain, had the most differences. 18323 has been shown by many typing methods to differ from most *B. pertussis* strains, lacking some genes usually seen in *B. pertussis* while having copies of genes normally found in *B. bronchiseptica* and *B. parapertussis_Hu_* but not *B. pertussis* [41–44]. These genomic differences are reflected in the phylogenetic tree, where 18323 is the closest *B. pertussis* strain to Complex IV **(S1 Figure)**. 18323 was the only *B. pertussis* strain containing *ctaD2*. 18323 also contained unique mutations, resulting in premature stop codons in *ccoP, cydB3, cyoB1,* and *cyoC1*, as well as losing *cyoA1* and *cyoD1* **(S2 Table).** Interestingly, 18323 was not the only *B. pertussis* strain that lost the ability to make Cyd3. Six of the 13 remaining Complex II strains contain a frameshift mutation in the same position as 18323 but are shifted into a different reading frame. The fact that these 6 strains all have identical mutations indicates that they may have evolved from a common ancestor with this mutation and that this mutation occurred after *B. pertussis* became its own subspecies.

Despite the differences within *B. pertussis* strains, a pattern emerges; excluding 18323, all these strains have maintained functional loci encoding Cyd1, Cta1, and Cyo1. Therefore, we hypothesize that these three cytochrome oxidases are necessary and sufficient for *Bordetella* infection within the mammalian respiratory tract, the environment in which *B. pertussis* has specialized, and the brief time spent between hosts during transmission.

### Cyd1, Cta1, or Cyo1 is required for growth in ambient air

To assess the necessity of each cytochrome oxidase for survival under various conditions, we generated *B. bronchiseptica* strains with in-frame gene loci deletions using allelic exchange. Starting with our wild-type strain RB50 (WT), we were able to generate strains lacking *cco* (Δ*cco*), *cyd1* (Δ*cyd1*), *cyd3* (Δ*cyd3*), *cio* (Δ*cio*), *cta1* (Δ*cta1*), *ctaD2* (Δ*ctaD2*), or *cyo1* (1′*cyo1*) **(S2 Figure)**. We were unable to delete the *cyo2* locus in its entirety; instead, we deleted *cyoABC2* and left *cyoD2* intact. The resulting strain lacks a functional complex generated by the proteins encoded by *cyo2* and is therefore called Δ*cyo2* hereafter.

Based on genome comparisons between *B. bronchiseptica* and *B. pertussis,* we aimed to generate strains that allowed us to test the necessity and sufficiency of the three cytochrome oxidase-encoding loci broadly conserved in *B. pertussis* (*cyd1, cta1,* and *cyo1*). We tried to make a strain that lacks *cyd1, cyo1*, and *cta1* but were unable to generate it using our normal growth conditions; all co-integrants reverted to WT at the last step of the allelic exchange procedure, suggesting that at least one of the *Bp*-conserved cytochrome oxidases is required for growth of *B. bronchiseptica* under standard laboratory conditions, i.e. in SS broth, in ambient air. We were able to make a strain with only the three conserved cytochrome oxidases by deleting the other five cytochrome oxidase-encoding loci (*cyd1^+^ cta1^+^ cyo1^+^,* abbreviated *Bp*-conserved). We then used this strain to generate strains for assessing the necessity of each of the cytochrome oxidases conserved in *B. pertussis* when only these three are present. We were able to create all three iterations; a strain with only *cta1* and *cyo1* (*cta1^+^ cyo1^+^*), a strain with only *cyd1* and *cyo1* (*cyd1^+^ cyo1^+^*), and a strain with only *cyd1* and *cta1* (*cyd1^+^ cta1^+^*).

To assess the sufficiency of each cytochrome oxidase for growth under various conditions, we generated strains that contain in-frame deletions of seven of the eight cytochrome oxidase-encoding loci. We were able to generate three of the eight possible strains with only one cytochrome oxidase-encoding gene loci: a strain with only *cyd1* (*cyd1^+^*), a strain with only *cta1* (*cta1^+^*), and a strain with only *cyo1* (*cyo1^+^*). We were unable to generate the five other possible strains with only one cytochrome oxidase-encoding gene locus remaining under our normal growth conditions; indeed, we were only able to generate strains lacking six of the eight oxidase-encoding gene loci if one of the two remaining loci was *cyd1, cta1,* or *cyo1* **(S2 Figure)**. This result, combined with the fact that we were unable to generate a strain missing all three cytochrome oxidases conserved in *B. pertussis,* indicates that Cyd1, Cta1, and Cyo1 are the primary cytochrome oxidases used under normal growth conditions in ambient air and that at least one is required under this condition.

### All of the strains we constructed are able to respire

To determine how well our strains were able to respire after losing one or more cytochrome oxidase(s), we grew our strains on medium containing triphenyl tetrazolium chloride (TTC). TTC accepts electrons from the electron transport chain. When TTC is reduced in the presence of cytochrome oxidases in a sufficiently dense culture, it undergoes an irreversible color shift from colorless to red [45]. We examined the color of colony biofilms, normalized by starting optical density (OD_600_), after 24 hours of growth on media containing TTC. All strains were able to respire, as evidenced by the red color of the colonies **(S3 Figure).** All strains except *cta1^+^*had a similar level of redness, indicating that the mutant strains had a similar level of respiration as WT. *cta1^+^* was less red overall but had spots with a higher density of red pigment than WT. This color variation could reflect heterogeneity in the mutant population.

*cta1^+^* has other phenotypes that differ from WT. This strain generated smaller colonies than WT on Bordet-Gengou (BG) blood agar plates, our standard culturing medium, requiring 72 hours to produce visible colonies as opposed to the 48 hours required by WT. Moreover, *cta1^+^* can’t grow in our normal liquid culturing medium, Stainer-Scholte (SS) broth, when taken from BG agar. However, colonies from BG agar are able to grow on SS agar, and these colonies can then be grown in SS broth. *cta1^+^* cultures grown in SS broth or on SS agar can grow on BG agar. This phenotype was not due to second-site mutations, as whole-genome sequencing revealed no unique mutations compared to WT, *cyd1^+^,* or *cyo1^+^*. The reason for this unusual phenotype is not known, and we did not pursue it further.

### No cytochrome oxidase is necessary for growth in ambient air, but Cyd1, Cta1, and Cyo1 are each sufficient

To determine if individual cytochrome oxidases are required for normal growth, we grew strains with deletions of individual cytochrome oxidase-encoding loci (i.e. 1′*cco,* 1′*cyd1,* Δ*cyd3,* 1′*cta1,* Δ*ctaD2,* 1′*cio,* 1′*cyo1,* and Δ*cyo2*) in SS medium in ambient air conditions and measured growth over time via OD_600_ and number of viable bacteria (CFU/mL). WT reached stationary phase between 12 and 18 hours (**Figure 1A**). While the OD_600_ of WT stayed consistent after the culture entered stationary phase, the CFU/mL began to decline after 30 hours. All eight strains lacking a single cytochrome oxidase phenocopied WT by both OD_600_ and CFU/mL (**Figure 1A**). This result indicates that no single cytochrome oxidase is required for growth under standard laboratory conditions.

**Figure 1.**
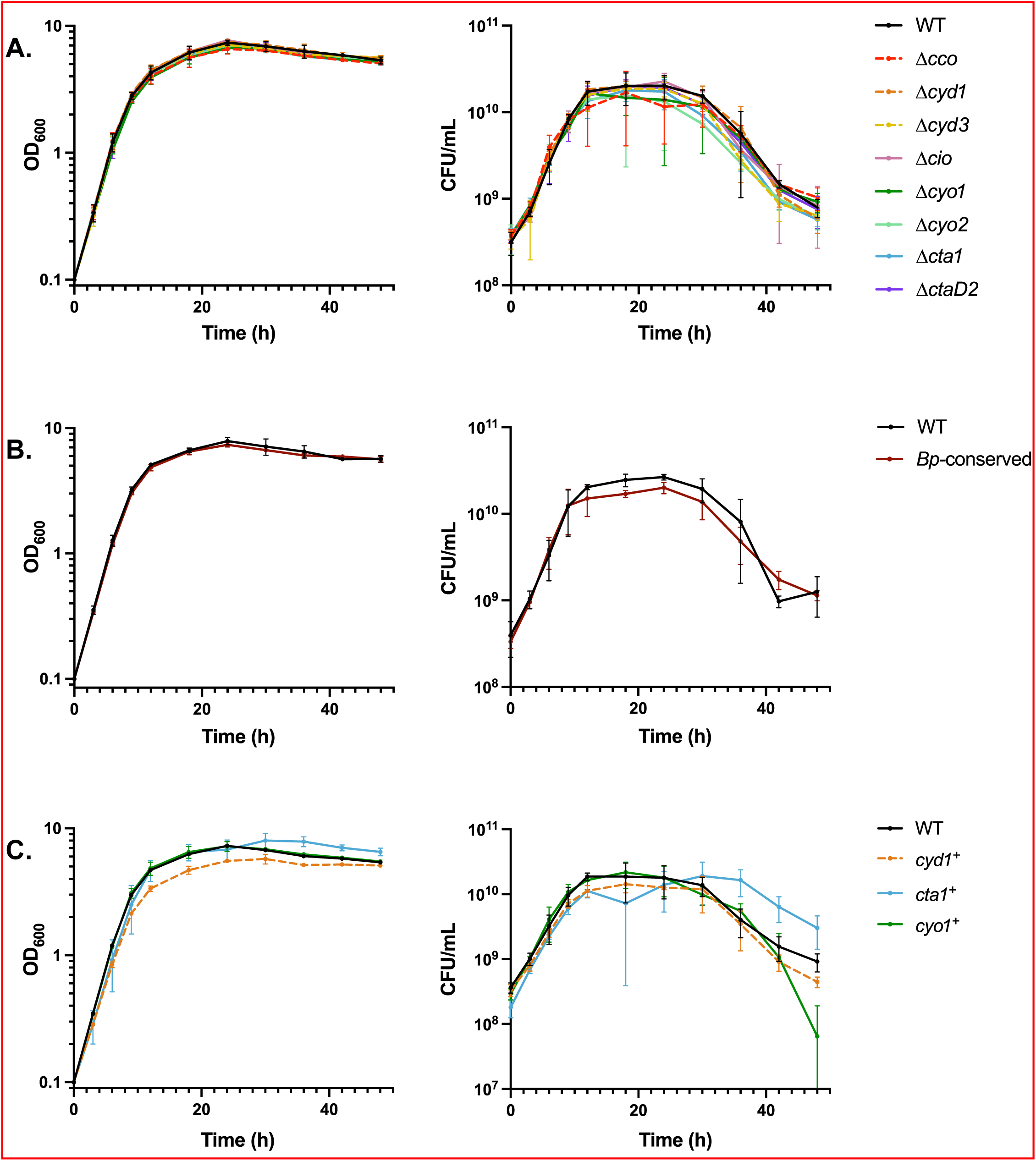
No cytochrome oxidase is necessary for growth in ambient air, but Cyd1, Cta1, and Cyo1 are each sufficient. Growth over time, measured via optical density (left) or CFU/mL (right), for strains lacking a single cytochrome oxidase-encoding gene loci (A), a strain with only the cytochrome oxidase-encoding gene loci conserved in *B. pertussis* (*Bp*-conserved) (B), or strains with only a single cytochrome oxidase-encoding gene loci (C). Cultures were started at 0.1 OD_600_, which contains approximately 3.5×10^8^ CFU/mL. Points represent the mean of at least 3 biological replicates, with bars representing the standard deviation.

To determine if the cytochrome oxidases broadly conserved in *B. pertussis* are sufficient for growth under the same conditions, we assessed growth of *Bp*-conserved. *Bp*-conserved phenocopied WT over time (**Figure 1B**), indicating that the combination of Cyd1, Cta1, and Cyo1 is sufficient under these conditions. Given that the combination of the three *Bp-*conserved cytochrome oxidases was sufficient, we then assessed their individual sufficiency. To do so, we used strains with only a single cytochrome oxidase-encoding locus remaining (i.e. *cyd1^+^, cta1^+^,* and *cyo1^+^*). Overall, these strains behaved similarly to WT, reaching stationary phase at the same time and having a similar growth rate (**Figure 1C**). *cyd1^+^* consistently had a slightly lower, but not significantly different, OD_600_ and CFU/mL than WT. After 30 hours, when WT began to decrease in CFU/mL, *cta1^+^* decreased in CFU/mL more slowly than WT, while *cyo1^+^* decreased in CFU/mL more quickly than WT, especially between 36 and 48 hours. These data indicate that each of the cytochrome oxidases conserved in *B. pertussis* is sufficient for WT levels of growth in liquid media in ambient air.

### None of the cytochrome oxidases conserved in *B. pertussis* are critical for growth in low oxygen or in high carbon dioxide

Under normal laboratory conditions, bacteria have access to ambient air, which contains approximately 21% O_2_. When growing within the mammalian respiratory tract, however, bacteria encounter a different atmospheric environment. Upon inhalation, alveoli inflate and oxygen in the air is exchanged with carbon dioxide (CO_2_) produced by cellular respiration; upon exhalation, this new gas mixture, which contains approximately 16% O_2_ and 5% CO_2_, is released back into the environment (reviewed in [46]). The classical bordetellae are likely further restricted from access to oxygen since the ciliated epithelial cells to which the bacteria adhere are coated in a layer of secreted mucus designed to protect the respiratory tract from environmental pathogens [47].

To investigate how different cytochrome oxidases contribute to the ability of *B. bronchiseptica* to grow in the different atmospheric conditions it may encounter in the mammalian respiratory tract, we grew strains within an incubator that allowed us to manipulate the atmospheric concentrations of oxygen, nitrogen, and CO_2_. We collected endpoint samples to keep the gas concentrations consistent for the duration of the experiment since taking samples required us to open the incubator, thus exposing the cultures to ambient air. Since we hypothesize that the three *Bp-*conserved cytochrome oxidases are sufficient for growth within the mammalian respiratory tract, we focused on assessing the growth of *Bp-*conserved, *cyd1^+^, cta1^+^,* and *cyo1^+^* relative to WT.

First, we assessed growth in low oxygen. Previous studies in *E. coli* have shown that atmospheric oxygen concentrations at or below 5% are sufficient to shift expression from low-affinity to high-affinity cytochrome oxidases in liquid cultures [48].We therefore grew our cultures under 5% O_2_ with constant agitation to maximize gas exchange. WT grew slower in 5% O_2_ than in ambient air; cultures grown for 48 hours at 5% O_2_ contained less than half the bacteria found at 24 hours in cultures grown in ambient air (**Figure 2A vs 2B).** Surprisingly, *Bp-*conserved, *cyd1^+^, cyo1^+^,* and *cta1^+^* were all able to grow to WT levels in 5% O_2,_ with no strain having a significant defect relative to WT in both OD_600_ and CFU/mL (**Figure 2B**). Since no cytochrome oxidase was critical for growth at 5% O_2_, we lowered the level of oxygen to 2% O_2_. Under 2% O_2,_ WT struggled to grow; cultures grown for 72 hours at 2% O_2_ contained approximately 13% of the bacteria found at 24 hours in cultures grown in ambient air (**Figure 2A vs 2C).** Despite low oxygen availability, *cyo1^+^* and *cta1^+^* were able to grow to similar levels as WT and *Bp-*conserved and *cyd1^+^* only had a slight growth defect (**Figure 2C**). Together, these data indicate that none of these cytochrome oxidases is required for growth under low oxygen conditions. Conversely, these data indicate that Cyd1, Cta1, or Cyo1 is sufficient for WT-levels of growth under low oxygen conditions.

**Figure 2.**
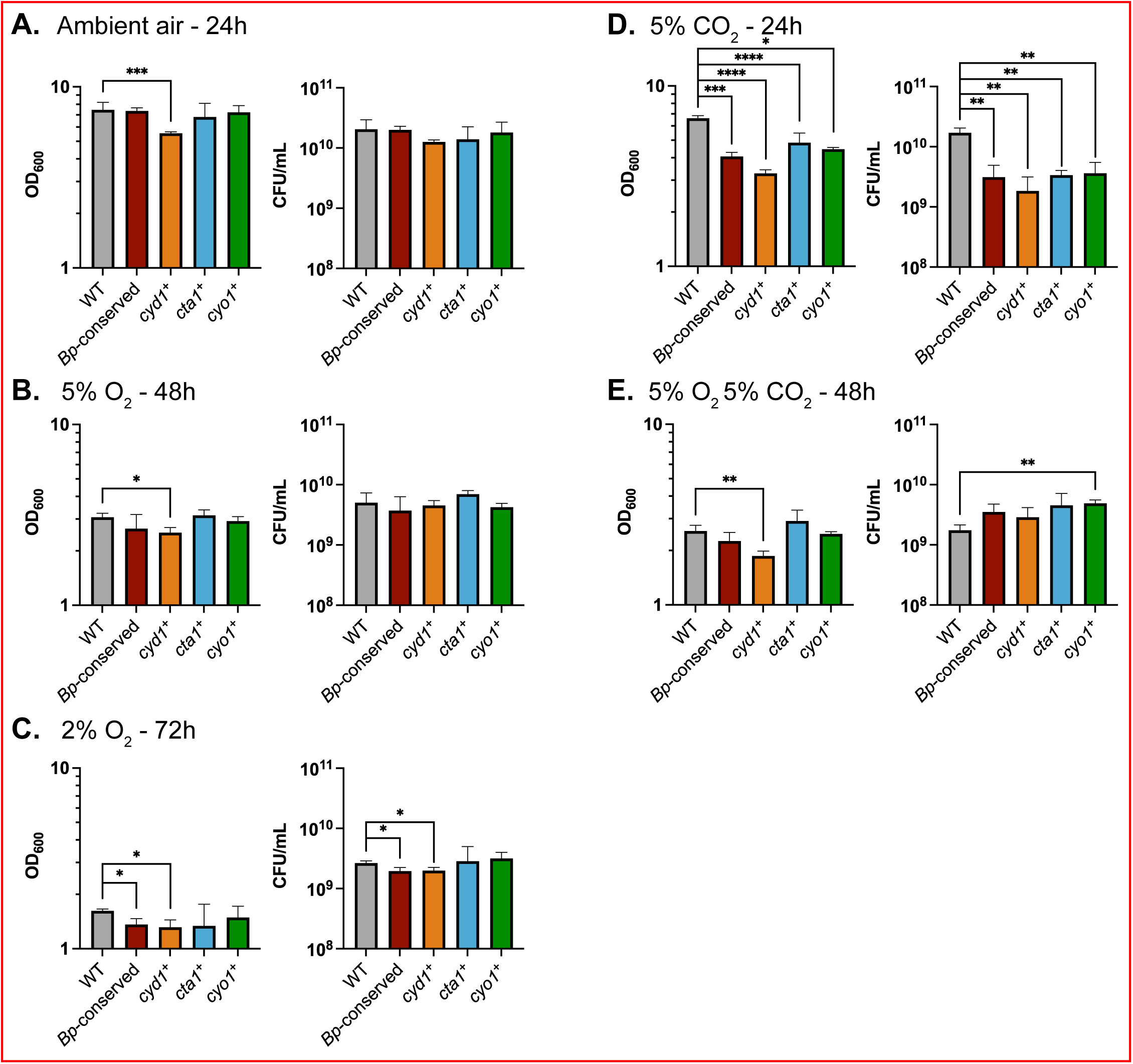
Growth under different atmospheric conditions relevant to the environment of the mammalian respiratory tract. Growth, measured via optical density (left) or CFU/mL (right), for RB50 (WT, gray), a strain with only the cytochrome oxidase-encoding gene loci conserved in *B. pertussis* (*Bp*-conserved, maroon), a strain with only *cydAB1* (*cyd1^+^,* orange), a strain with only *ctaCDFGE1* (*cta1^+^*, blue), or a strain with only *cyoABCD1* (*cyo1^+^*, green). Cultures were started at 0.1 OD_600_, which contains approximately 3.5×10^8^ CFU/mL. Cultures were grown in either: ambient air for 24 hours (A), 5% O_2_ for 48 hours (B), 2% O_2_ for 72 hours (C), 5% CO_2_ for 24 hours (D), or 5% O_2_ 5% CO_2_ for 48 hours (E). The data from A are identical to the 24-hour timepoint of Figure 1A and are included for comparison. Bars represent the mean of at least 3 biological replicates, with error bars representing the standard deviation. Statistical significance was determined using unpaired Student’s t-test. *, p < 0.05; **, p < 0.005; ***, p < 0.0005; ****, p < 0.0001.

Since bacteria in the respiratory tract also encounter increased carbon dioxide, we next grew our strains in 5% CO_2_ to determine if growth is impacted by this change. WT grew similarly in 5% CO_2_ as in ambient air (**Figure 2A vs 2D).** *Bp-*conserved, *cyd1^+^, cyo1^+^,* and *cta1^+^,* however, were all defective relative to WT by both OD_600_ and CFU/mL (**Figure 2D**). This result is surprising, as it suggests that the three cytochrome oxidases conserved in *B. pertussis* are not sufficient for efficient growth in the presence of 5% CO_2_. Indeed, it suggests that at least one of the cytochrome oxidases deleted in *Bp-*conserved contributes to growth in 5% CO_2_.

Given that the respiratory environment can simultaneously contain higher levels of carbon dioxide and lower levels of oxygen than ambient air, we tested how well our strains are able to grow in 5% O_2_ 5% CO_2_. Under this condition, WT grew similarly to how it grew in 5% O_2_ alone (**Figure 2B vs 2E).** After 48 hours of growth, *Bp-*conserved, *cyd1^+^, cyo1^+^,* and *cta1^+^* were all able to grow to WT levels, with no strain having a significant defect relative to WT by both OD_600_ and CFU/mL (**Figure 2E**).

### No single cytochrome oxidase is required for persistence in the murine respiratory tract

To determine the role of individual cytochrome oxidases during infection, we inoculated mice intranasally with bacteria using our high-dose, large-volume protocol, which allows us to assess how well different strains can persist once introduced in the respiratory tract. We first used strains containing deletions of individual cytochrome oxidase-encoding loci to determine which cytochrome oxidases, if any, are necessary for persistence. We then collected the right lung, the trachea, and nasal cavity tissue at 3 hours, 1 day, 3 days, and 7 days post-inoculation and determined the bacterial burden in each organ. Due to the number of strains involved, we could not test all eight strains simultaneously. Instead, we tested them in batches, with each including WT, then compiled the data. All strains caused the same bacterial burden in mice as WT in the nasal cavity, trachea, and lung (**Figure 3**). These data indicate that no individual cytochrome oxidase is required for persistence in the murine respiratory tract.

**Figure 3.**
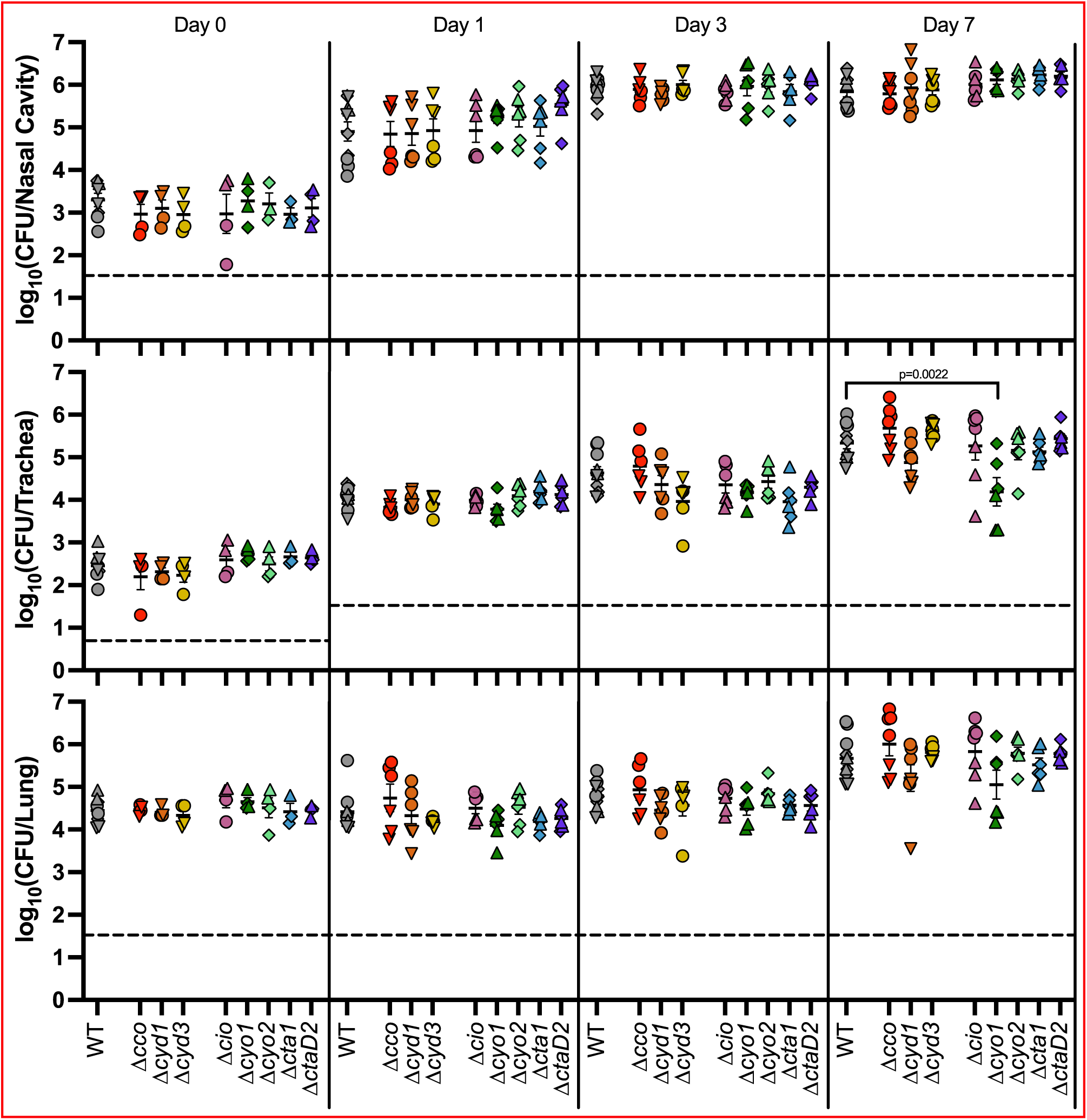
No single cytochrome oxidase is required during murine infection. Bacterial burden over time within the nasal cavity (upper), trachea (middle), and right lung (lower) of mice infected with either RB50 (WT) or strains lacking a single cytochrome oxidase-encoding gene loci. Different symbols represent different batches of inoculations, which always include WT for comparison: circles (1′*cco,* 1′*cyd1,* 1′*cyd3*, and 1′*cio*), upright triangles (1′*cio,* 1′*cyo1,* 1′*cyo2,* 1′*cta1,* and 1′*ctaD2*), upside-down triangles (1′*cco,* 1′*cyd1*, and 1′*cyd3*), and diamonds (1′*cyo1*, 1′*cyo2,* 1′*cta1*, and 1′*ctaD2*). Each point represents data from one mouse. n=4 for day 0, n=6 for all other timepoints, from two independent experiments, for all mutant strains. Each point represents a single mouse. Dashed line represents the limit of detection. Statistical significance was determined using unpaired Student’s t-test; p-values are indicated when p<0.01.

### The cytochrome oxidases conserved in *B. pertussis* are sufficient for persistence during murine infection

We next assessed the role of the three cytochrome oxidases conserved in *B. pertussis* for persistence during infection. We infected mice with either *Bp*-conserved or WT, and determined the bacterial burden in the nasal cavity, trachea, and lung over the course of infection. The *Bp-*conserved strain was statistically indistinguishable from WT in all tissues at all time points tested (**Figure 4**), indicating that the three cytochrome oxidases conserved in *B. pertussis* are sufficient for persistence in mice.

**Figure 4.**
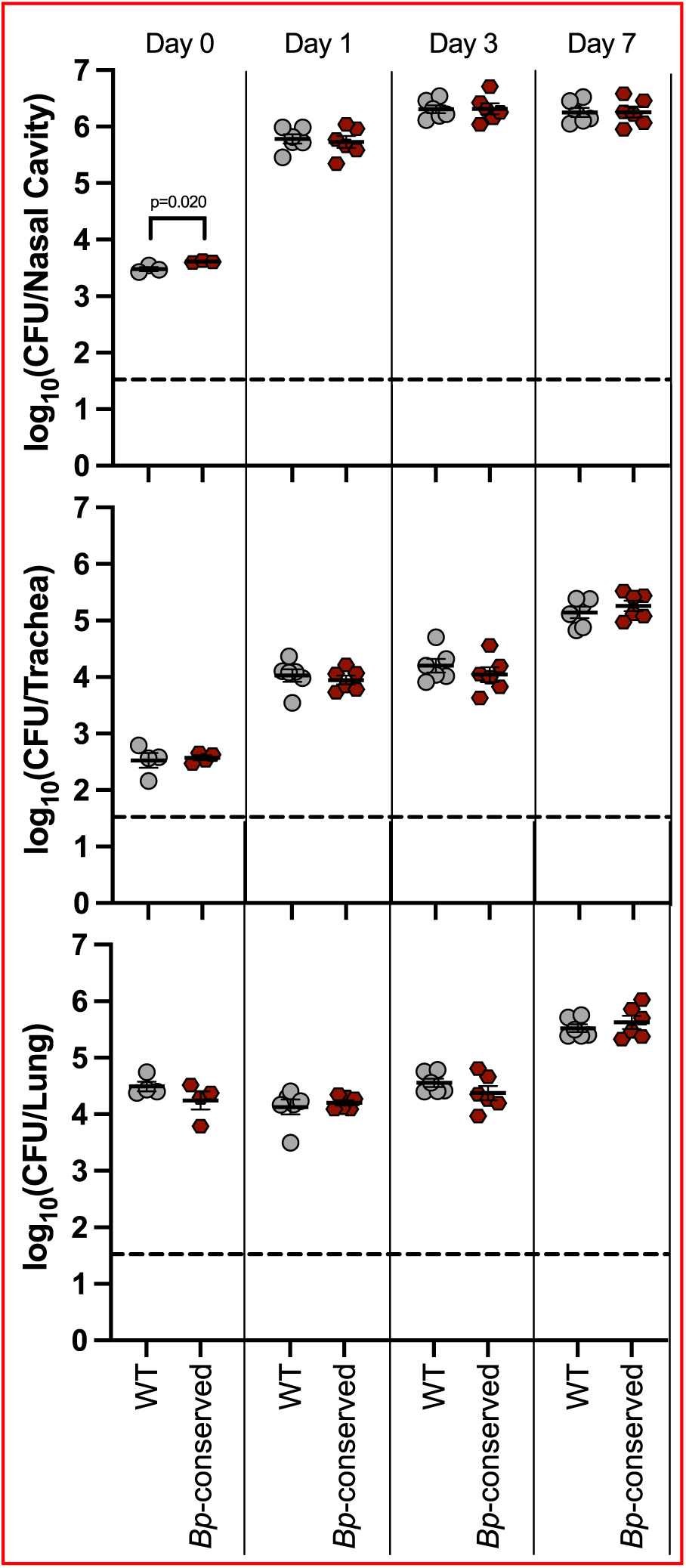
The three cytochrome oxidases conserved in *B. pertussis* are sufficient for persistence in the murine respiratory tract. Bacterial burden over time within the nasal cavity (upper), trachea (middle), and right lung (lower) of mice infected with either wild-type bacteria (WT, grey circles) or a strain with only the cytochrome oxidase-encoding gene loci conserved in *B. pertussis* (*Bp*-conserved, maroon hexagon). n=4 for day 0, n=6 for all other timepoints, with each point representing a single mouse. Samples were collected from two independent experiments. Dashed line represents the limit of detection. Statistical significance was determined using unpaired Student’s t-test; p-values are indicated when p < 0.05.

### Cyo1 is sufficient for persistence during infection, while strains with only Cyd1 or Cta1 are defective for persistence relative to WT

The three cytochrome oxidases conserved in *B. pertussis* are predicted to have different characteristics (**Table 1**). Cyd1 is predicted to be a high-affinity *bd-*type ubiquinol oxidase. Cta1 is predicted to be a low-affinity HCO cytochrome *c* oxidase. Cyo1 is predicted to be a low-affinity HCO ubiquinol oxidase. We hypothesize that each of these cytochrome oxidases fulfills a different function during infection due to their different biochemical properties. To test this hypothesis, we infected mice using strains with only a single cytochrome oxidase-encoding gene locus remaining: *cyd1^+^, cta1^+^,* and *cyo1^+^*. *cyd1^+^* had a slight defect relative to WT in the nasal cavity, causing a lower bacterial burden at day 1 that was able to reach WT levels by day 3 (**Figure 5A, top)**. *cyd1^+^* was consistently defective in the lower respiratory tract relative to WT, causing a significantly lower bacterial burden in the trachea and the lung out to 14 days (**Figure 5A, middle and bottom)**. *cta1^+^* was defective relative to WT in all three sampled tissues (**Figure 5B**). However, this strain was not cleared from the lung faster than WT, as seen by the statistically similar burdens recovered on day 14 when the burden of WT decreased relative to earlier time points (**Figure 5B, bottom)**. Most surprisingly, *cyo1^+^* was recovered at levels similar to WT in all three sampled tissues across the course of infection (**Figure 5C**). These data indicate that Cyo1 is sufficient for WT-levels of growth in this model of mouse infection, despite being predicted to be optimized for ambient air levels of oxygen, while Cyd1 and Cta1 are not sufficient. Additionally, all three strains are able to persist to some degree throughout the respiratory tract once introduced, indicating that any of these cytochrome oxidases is sufficient for survival within a mouse.

**Figure 5.**
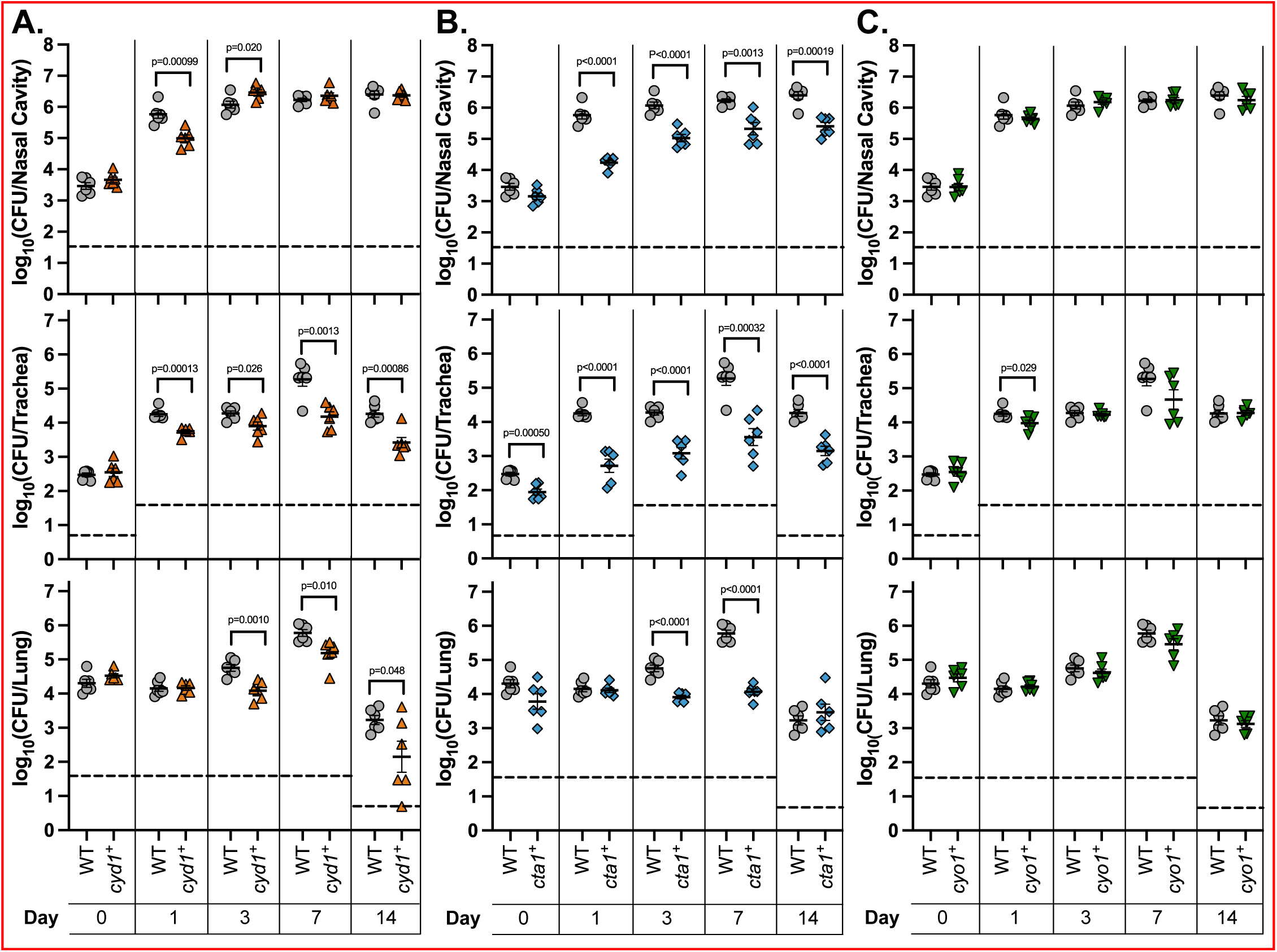
Cyo1 is sufficient for WT-levels of persistence within the mammalian respiratory tract, while Cyd1 and Cta1 are not. Bacterial burden over time within the nasal cavity (upper), trachea (middle), and right lung (lower) of mice infected with wild-type bacteria (WT, grey circles), a strain with only *cydAB1* (*cyd1^+^,* orange upright triangles), a strain with only *ctaCDFGE1* (*cta1^+^*, blue diamonds), or a strain with only *cyoABCD1* (*cyo1^+^*, green upside-down triangles). These strains were tested in the same experiment but separated into separate graphs for analysis; thus, the mutant strains are compared to the same WT data. n=4 for day 0, n=6 for all other timepoints, with each point representing a single mouse. Samples were collected from two independent experiments. Dashed line represents the limit of detection. Statistical significance was determined using unpaired Student’s t-test; p-values are indicated when p < 0.05.

### The *Bp-*conserved strain is indistinguishable from WT in its ability to establish respiratory colonization

We and others have used high-dose, large-volume and low-dose, small-volume mouse protocols to investigate respiratory tract persistence and establishment of infection, respectively. In our lab, we have previously used the low-dose, small-volume protocol only in rats, where we showed that doses of 20 CFU are sufficient for infection in rats [49]. To adapt this approach to mice, we performed preliminary experiments using WT to determine a viable volume and dose for intranasal inoculation in wild-type mice. We found that 4μL was sufficient to introduce bacteria to the nasal cavity but not the trachea, as we were able to recover CFU from the nasal cavity but not the trachea at this volume on the same day as we inoculated the mice **(S4A Figure).** We also found that 100 CFU per inoculum was sufficient to ensure robust colonization of the nasal cavity, as well as consistently recoverable burden from the trachea by day 7 post-inoculation **(S4A Figure).** In this model, bacteria were not consistently recovered from the lungs.

We next assessed whether the three *Bp-*conserved cytochrome oxidases are sufficient for infection. *Bp-*conserved phenocopied WT in the high-dose, large-volume model (**Figure 4),** demonstrating that the three cytochrome oxidases found in *B. pertussis* are sufficient for persistence. We performed a preliminary experiment to determine if these three cytochrome oxidases are also sufficient for establishing infection. Mice infected with *Bp-*conserved had a similar bacterial burden in the nasal cavity 3 days post-inoculation as mice infected with WT, indicating that the combination of Cyd1, Cyo1, and Cta1 is sufficient for establishing infection **(S3B Figure).**

### Cyo1 is sufficient for efficient establishment of infection, while Cyd1 and Cta1 are not

To assess the individual roles of Cyd1, Cyo1, and Cta1 during the establishment of infection, we first tested whether any of these three cytochrome oxidases is required. We used the derivatives of *Bp-conserved* that have two remaining cytochrome oxidases (i.e. *cta1^+^ cyo1^+^, cyd1^+^ cyo1^+^,* and *cyd1^+^ cta1^+^*). These strains phenocopied WT growth in ambient air **(S5A Figure)**. We infected mice with these strains using our low-dose, small-volume protocol and determined the nasal cavity bacterial burdens 3 and 7 days post-inoculation. We found that mice infected with the strains with two cytochrome oxidases had similar bacterial burdens to mice infected with *Bp-*conserved **(S5B Figure)**. One strain, *cyd1^+^ cta1^+^,* trended towards causing a lower bacterial burden than *Bp-*conserved, but this difference was not significant. Based on these data, no single cytochrome oxidase is required to establish infection.

To determine if any of the three *Bp*-conserved cytochrome oxidases is sufficient to establish infection, we infected mice with the strains with only a single cytochrome oxidase-encoding locus remaining (i.e. *cyd1^+^, cta1^+^,* and *cyo1^+^*) and determined the bacterial burden as above. Overall, the trends in the low-dose, small-volume experiments mirrored those seen in the high-dose, large-volume experiments. Mice infected with *cyd1^+^* had a lower bacterial burden 3 days post-inoculation than mice infected with *Bp-*conserved but reached the same burden by day 7 (**Figure 6**). Mice infected with *cta1^+^*had a significantly lower bacterial burden than mice infected with *Bp-*conserved at both day 3 and 7 (**Figure 6**). Despite causing less than one percent of the bacterial burden of *Bp-*conserved strain at day 7 in mice, *cta1^+^*nonetheless was able to establish infection and persist for the duration of the experiment. Mice infected with *cyo1^+^* had the same bacterial burden as mice infected with *Bp-*conserved at both timepoints (**Figure 6**). Together, these data indicate that while all three cytochrome oxidases are sufficient to enable the establishment of infection, as seen by recoverable bacteria following infection with all three strains, only Cyo1 is sufficient to achieve the level of colonization reached when all three cytochrome oxidases conserved in *B. pertussis* are present.

**Figure 6.**
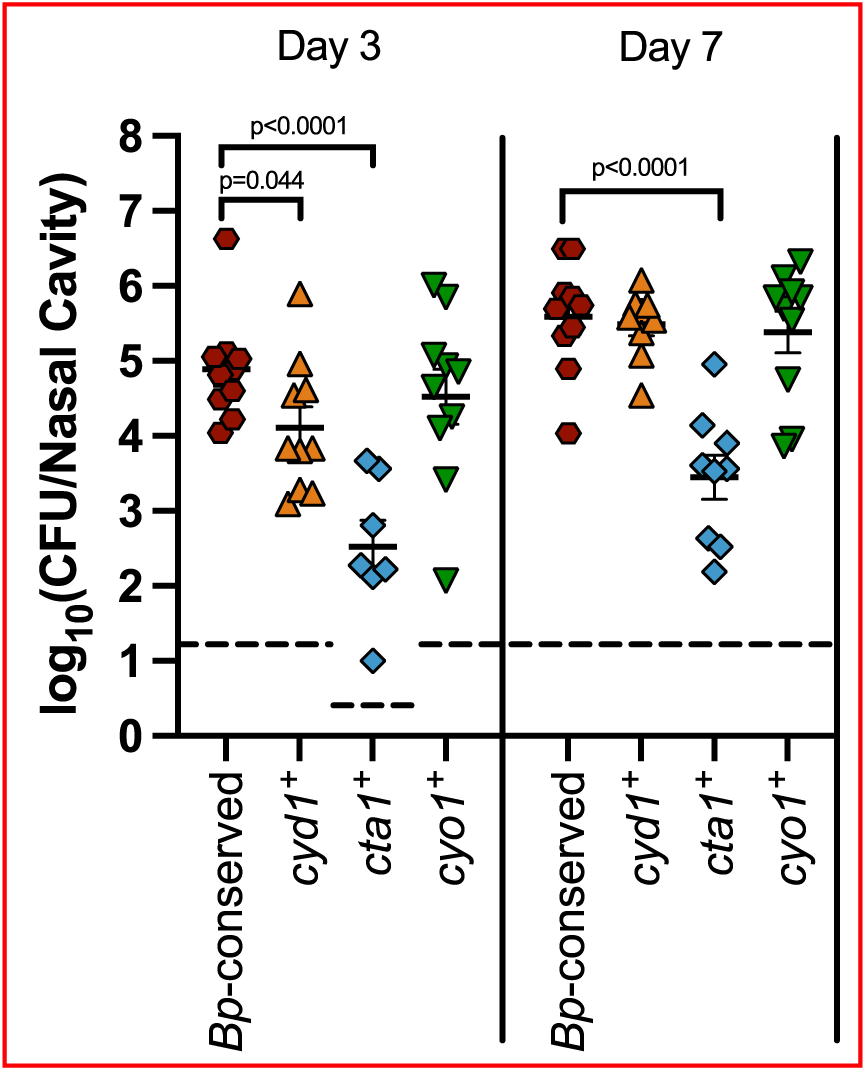
Cyo1 is sufficient for the efficient establishment of colonization, while Cyd1 and Cta1 are not. Bacterial burden over time within the nasal cavity of mice infected with a strain with only the cytochrome oxidase-encoding gene loci conserved in *B. pertussis* (*Bp*-conserved, maroon hexagon), a strain with only *cydAB1* (*cyd1^+^,* orange upright triangles), a strain with only *ctaCDFE1* (*cta1^+^*, blue diamonds), or a strain with only *cyoABCD1* (*cyo1^+^*, green upside-down triangles). Samples from 10 mice across two independent experiments were collected at each timepoint for each strain. However, due to the natural microbiota of the nasal cavity, *B. bronchiseptica* could not always be innumerated due to contamination. Therefore, n=7 for *cta1^+^* day 3, n=9 for *cyd1^+^*day 7 and *cta1^+^* day 7. Each point represents a single mouse. Dashed line represents the limit of detection. Statistical significance was determined using unpaired Student’s t-test; p-values are indicated when p < 0.05.

### Regulation of cytochrome oxidase-encoding genes broadly follows the predicted affinity of the cytochrome oxidases they encode

Given that we were unable to construct a strain that lacks all three of the cytochrome oxidase-encoding gene loci conserved in *B. pertussis* **(S2 Figure)**, we investigated how different cytochrome oxidase-encoding genes are regulated, hypothesizing that some loci may not be expressed under laboratory growth conditions. We diluted overnight cultures to the same density, grew them for 4 hours in either ambient air, 5% O_2_, 2% O_2,_ or 5% CO_2_, extracted RNA, and sent it for sequence analysis.

First, we examined the level of transcripts in ambient air to investigate the expression of cytochrome oxidase-encoding genes under our normal growth conditions. Raw normalized read counts were converted to counts per million for comparison. All eight cytochrome oxidase-encoding gene loci had detectable transcripts (**Figure 7A**). However, when compared to *fhaB* and *flaA,* two well-characterized and highly-regulated genes, we found that transcript levels for most of the non-conserved cytochrome oxidase-encoding gene loci (*cyd3, cio, ctaD2,* and *cyo2*) were similar to that of *flaA,* the protein product of which cannot be detected under these growth conditions [50]. Therefore, transcription of *cyd3, cio, ctaD2,* and *cyo2* may be insufficient to produce enough functional enzyme complexes to support respiration. By contrast, transcript abundance for *ccoN* was the highest of all of the loci, with high expression of the rest of the *cco* locus as well, yet we were unable to make a strain with only *cco*. This result suggests a lack of sufficient translation, complex assembly, or cofactors required for Cco. The three *B. pertussis*-conserved cytochrome oxidase-encoding loci had high levels of expression, though not as high as *fhaB,* which encodes a highly abundant secreted protein.

**Figure 7.**
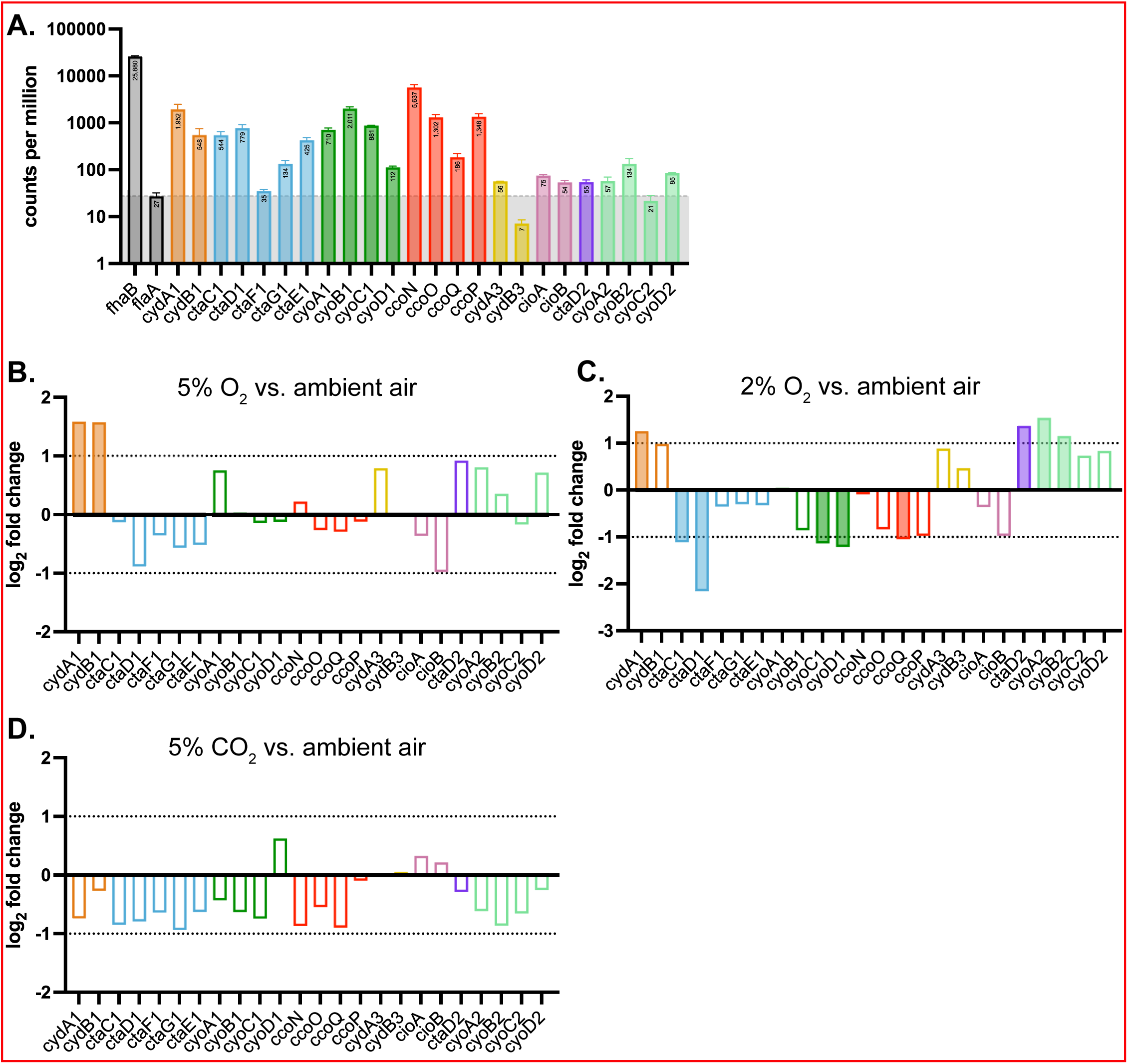
Regulation of cytochrome oxidase genes under different atmospheric conditions, measured using RNA sequencing. (A) Transcript abundance, measured by counts per million, for cytochrome oxidase-encoding genes from *B. bronchiseptica* grown in ambient air. *fhaB* and *flaA*, two highly-regulated and well-characterized genes, are included for comparison. *flaA* transcription level (gray shading) are used as a reference, as *flaA* is negatively regulated under this condition and no protein product can be detected. Numbers in the bars represent the mean number of transcripts. (B-D) Comparative transcription, measured by fold change in transcript abundance, comparing 5% O_2_ (B), 2% O_2_ (C), or 5% CO_2_ (D) to ambient air. Genes with log_2_ fold changes greater than 1 or less than −1 have filled-in bars. Differential expression analysis was performed using edgeR’s glmQLFTest.

We next examined the fold change in transcript abundance in bacteria grown in our conditions of interest (5% O_2_, 2% O_2_, and 5% CO_2_) and ambient air. In 5% O_2_, *cydA1* and *cydB1* had ∼3 times more transcripts than in ambient air (**Figure 7B**). This result is consistent with the prediction that Cyd1 functions as a high-affinity cytochrome oxidase and is therefore adapted for use in low oxygen environments. No other gene had a significant difference in transcript abundance in 5% O_2_ versus ambient air.

In 2% O_2_, differences in transcript abundance were greater (**Figure 7C**). *cydA1* had significantly more transcripts in 2% O_2_ than in ambient air but fewer transcripts than in 5% O_2_. Genes within *cyo1* (*cyoC1* and *cyoD1*) had significantly fewer transcripts in 2% O_2_ relative to ambient air, which matches the prediction that these genes encode components of a low-affinity cytochrome oxidase. *ctaC1* and *ctaD1* have less expression in 2% O_2_ relative to ambient air, with *ctaD1* having a larger relative decrease in transcripts. Interestingly, *ctaD2* had higher expression in 2% O_2_ relative to ambient air, with ∼2.6 times more transcripts in ambient air than in 2% O_2_. Since the genes encoding other components of Cta1 did not significantly decrease in expression in 2% O_2_, CtaD2 could be interacting with CtaC1 and CtaE1 (and potentially CtaF and CtaG1) to form a heterocomplex (i.e. CtaC1<u>D2</u>F1G1E1). Unexpectedly, *ccoQ* had significantly lower levels of transcripts in 2% O_2_ relative to ambient air, despite being predicted to encode part of a high-affinity cytochrome oxidase, and *cyoC2* and *cyoD2* had significantly higher levels of transcripts, despite being predicted to encode parts of a low-affinity cytochrome oxidase.

In 5% CO_2_, no gene had significant changes in expression relative to growth in ambient air (**Figure 7D**). This was surprising, as 5% CO_2_ was the only growth condition tested that resulted in a large growth defect in *Bp*-conserved relative to WT (**Figure 2**), leading us to hypothesize that a non-*Bp-*conserved cytochrome oxidase that is not required for growth in ambient air must be important for growth in 5% CO_2_. While this hypothesis could still be true, these results indicate that differences in growth in ambient air and 5% CO_2_ are not due to differences in cytochrome oxidase-encoding gene expression.

## DISCUSSION

Our goal was to understand the importance of cytochrome oxidases for bacterial pathogens that infect the mammalian respiratory tract. *B. bronchiseptica* contains eight cytochrome oxidase-encoding gene loci, while *B. pertussis,* which is strictly a human pathogen that survives only briefly in the environment during transmission between hosts, contains only three cytochrome oxidase-encoding gene loci that are broadly conserved across strains. Perhaps unsurprisingly, the three *B. pertussis*-conserved cytochrome oxidases were sufficient for *B. bronchiseptica* to both establish infection and persist in the respiratory tracts of mice. Somewhat surprisingly, however, *B. bronchiseptica* producing only a single low-affinity cytochrome oxidase (encoded by *cyoABCD1,* abbreviated Cyo1) was indistinguishable from WT in both infection models.

While it is well known that we inhale ambient air containing ∼21% O_2_ and low levels of CO_2_ and exhale air containing ∼16% O_2_ and ∼5% CO_2_, the levels of oxygen within specific microenvironments in the respiratory tract are unknown. The classical bordetellae bind to the surface of ciliated epithelial cells lining the respiratory tract, which do not continue into the smallest bronchioles or the alveoli. The ciliated epithelium is covered in a layer of mucus secreted by neighboring goblet cells and tethered mucins produced by the ciliated cells themselves. These viscous layers protect the respiratory tract from environmental pathogens but impede the diffusion of molecules, including oxygen, to the epithelium. We therefore anticipated that a high-affinity cytochrome oxidase would be important in this environment, and hence were surprised that *cyo1^+^*, a strain producing only a predicted low-affinity cytochrome oxidase, was able to infect mice with the same efficiency as WT. If the Cyo1 enzyme is, in fact, low-affinity as predicted, then this result indicates that the surface of the ciliated epithelium has access to more oxygen than previously thought.

The strain producing only Cyd1, by contrast, was attenuated relative to WT in both animal models. The level of attenuation depended on the site of the respiratory tract examined. In the nasal cavity, the *cyd1^+^* strain caused lower bacterial burdens than WT early in infection, but was able to reach WT-levels later in infection, indicating a growth delay in this environment. In the lower respiratory tract, however, the *cyd1^+^* strain was consistently defective relative to WT throughout the course of infection. If the surface of the ciliated epithelium is sufficiently oxygenated, as the results with the *cyo1^+^* strain suggest, this result could reflect the lower efficiency of high-affinity cytochrome oxidases, which pump fewer protons, and therefore generate less proton motive force, per electron transferred [6]. *bd-*type cytochrome oxidases have been linked to virulence in other pathogens not only for their oxygen affinity but also for their unique resiliency in the face of immune system attacks. *bd-*type cytochrome oxidases are more tolerant of oxidative and nitrosative stress than HCOs and have been shown to detoxify hydrogen peroxide and peroxynitrite produced by neutrophils and macrophages [51]. This role for *bd-*type cytochrome oxidases has been shown in *M. tuberculosis*, where the *bd-*type cytochrome oxidase is dispensable for the establishment of infection but is necessary after the initiation of adaptive immunity [18]. In this way, Cyd1 may play a larger role during in *B. pertussis,* which causes an acute infection that is combated by a strong, and ultimately sterilizing, mucosal immune response, compared to *B. bronchiseptica,* which typically causes chronic and, in the case of our mouse models, asymptomatic infection (reviewed in [52,53]).

Cta1 was the least effective during infection, with the strain producing only this cytochrome oxidase being recovered at lower levels than WT in all tissues at all timepoints in both models of infection. Cta1 could, however, play a role that was not assessed by our models. Indeed, previous microarray data indicated that *ctaC1, ctaD1, ctaG1,* and *ctaE1* are repressed by BvgAS, the two-component system that positively regulates all known protein virulence factors in *Bordetella* [54]. Given that the Bvg^+^ mode is both necessary and sufficient for infection, Cta1 may play a role outside the host during transmission [55].

Importantly, the mouse respiratory tract is not identical to the respiratory tract of humans; it is more similar to the human distal airways, both in terms of surface area and mucus composition. In humans, proximal airways are coated in mucus primarily made of MUC5AC mucins and distal airways have mucus made of MUC5B, whereas in mice, Muc5b is constitutively produced throughout the airway and almost no Muc5ac is produced [56]. The organization of the mucus is also different, with humans and other large mammals producing long thick bundles of mucins such that the proximal airway is coated in a layer of mucus, while in mice (and most likely, the distal airways of humans), the mucus is unevenly distributed, forming a patchy covering [57,58]. This patchy covering creates areas with thinner mucus layers, which would make oxygen diffusion easier. Therefore, the bacteria attached to the mouse ciliated epithelium may have access to more oxygen than they would on the human ciliated epithelium.

It is perhaps unsurprising that *B. bronchiseptica* can grow in low oxygen given its many cytochrome oxidase-encoding loci, three of which are predicted to encode high-affinity cytochrome oxidases. WT was able to grow in both 5% and 2% O_2_ (albeit at a far slower rate than in ambient air). It was surprising to us, however, that the *Bp*-conserved, *cyd1^+^, cta1^+^,* and *cyo1^+^* strains were all able to grow similarly under low oxygen conditions. Indeed, the *cta1^+^* and *cyo1^+^* strains, which encode only low-affinity cytochrome oxidases, grew as well as WT in 2% O_2_. This result indicates that all of the *B. pertussis-*conserved cytochrome oxidases all function equally well under low oxygen in laboratory conditions, despite their predicted affinities. Additionally, *B. bronchiseptica* is resilient when challenged with low oxygen. It is able to persist in static PBS, where oxygen diffusion is limited, with viable CFU being recovered even 24 weeks post-inoculation [59]. In this aspect, *B. bronchiseptica* is comparable to *M. tuberculosis,* another obligate aerobe that can persist in low oxygen conditions for long periods of time [60].

Our analysis primarily focused on the three cytochrome oxidase-encoding loci conserved in *B. pertussis.* However, we also examined the expression of the other five loci to see if they played a role in any of our experimental conditions. We found that *cyd3, cio, ctaD2,* and *cyo2* had fewer transcripts per million than *flaA*, a BvgAS-repressed gene whose protein product is not detected in our growth condition (SS medium) [50]. We were also unable to generate strains with only *cco, cyd3, cio, ctaD2,* or *cyo2* intact; at least one of the three *B. pertussis*-conserved loci had to be intact to generate a strain. Although the specific environments that *B. bronchiseptica* occupies outside the mammalian respiratory tract are unknown, our results suggests these environments are complex and varied, requiring different cytochrome oxidases than those utilized in lab culture.

In our RNAseq analyses, we only examined expression in wild-type *B. bronchiseptica* and therefore did not assess the effect that altering cell homeostasis by deleting cytochrome oxidase-encoding genes could have on global gene expression. In the *cyd1^+^, cta1^+^,* and *cyo1^+^*strains, it is likely that the remaining cytochrome oxidase-encoding locus is more highly expressed to compensate for the loss of the other two loci, which are normally expressed in our growth conditions. Expression changes could also explain the unusual plate-based growth phenotypes we observed for the *cta1^+^* strain, which could not be explained by secondary mutations. Upregulation of the remaining cytochrome oxidase-encoding loci in the strains with only a single locus could be masking synergy that occurs when the cytochrome oxidases are present at their endogenous levels.

Two additional trends emerged from the RNAseq. First, many highly BvgAS-activated genes had lower expression in low oxygen than in ambient air, including *bvgA*, *bvgS*, *fhaB*, and *cyaA* **(S3,4 Table).** By contrast, the common reference genes *rpoD* and *recA* were not differentially expressed in low oxygen as compared to ambient air, indicating that the differences seen for BvgAS-activated genes are not due to global changes in gene expression. *In vitro* studies using purified solubilized BvgS have shown that BvgS activity is sensitive to the redox state of ubiquinone through its PAS domain [61]. *In vivo* studies also support that BvgS is sensitive to redox potential via its PAS domain [62]. Thus, decreased expression of BvgAS-activated genes could be directly related to decreased respiration affecting BvgS activity in low oxygen environments.

Second, we found that expression of the *ptx* genes (*ptxABCDE*), which encode pertussis toxin, was increased in *B. bronchiseptica* in low oxygen **(S3,4 Table)**. This result is exciting, because *ptx* gene expression have never been detected in *B. bronchiseptica in vitro* or *in vivo,* and only a weak luminescent signal was detected when the *B. bronchiseptica ptx* promoter was used to drive the expression of luciferase-encoding genes [63]. Pertussis toxin is an important virulence factor during human infection, and in *B. pertussis,* the *ptx* genes are activated by BvgAS (reviewed in [64]). In Complex I *B. bronchiseptica* strains, from which the other classical bordetellae evolved, the *ptx* promoter region is not BvgAS-regulated but the *ptx* genes (as well as the *ptl* genes that encode the export system for pertussis toxin) are present and intact [65,66], suggesting that in *B. bronchiseptica*, pertussis toxin plays a different role than contributing to mammalian infection. By contrast, many Complex IV *B. bronchiseptica* strains have lost their *ptx* genes [37]. These strains also don’t survive as well as Complex I strains when grown in environmental conditions, instead being more adapted to the human hosts from which they were isolated. These results combined, together with the fact that the substrate for pertussis toxin is a eukaryotic G protein, suggest that pertussis toxin was used by ancestral *B. bronchiseptica* strains to manipulate a non-mammalian eukaryotic host or predator in the environment. As strains evolved to infect humans and lost their ability to survive in the environment, those that became *B. pertussis* acquired mutations in the *ptx* promoter that allowed for activation by BvgAS (and presumably not the regulator(s) that control *ptx* expression in *B. bronchiseptica*). Our findings suggest that one feature of the environment in which the *ptx* genes are expressed in *B. bronchiseptica* is low levels of accessible oxygen.

Previous genomic and experimental comparisons between the classical bordetellae have focused on virulence-associated genes like *fhaB, cyaA,* and the *ptx* genes. By comparing genes encoding proteins important for bacterial physiology, like cytochrome oxidases, we can also learn about which bacterial processes and pathways are important for survival within the host. Comparing *B. pertussis* and *B. bronchiseptica* led to our hypothesis that the three cytochrome-oxidase encoding gene loci that are conserved in *B. pertussis* would be sufficient during infection, which was supported by our experimental findings. Conversely, we can also learn about what is important for surviving in the environment. HT200, a *B. bronchiseptica* environmental isolate, differs from the other examined *B. bronchiseptica* strains in that HT200 does not have intact copies of all eight cytochrome-oxidase encoding gene loci. In particular, this strain has premature stop codons in both *cyoB1* and *cyoB2*, meaning it likely has no functional *bo_3_-*type cytochrome oxidases (Table S2). This result, especially in contrast with the broad conservation of *cyo1* within *B. pertussis* strains, indicates that while Cyo1 is important for survival within mammalian hosts, it is not important for survival within aqueous environments. This result also supports the hypothesis that the many different cytochrome oxidases encoded within the *B. bronchiseptica* genome all contribute to the ability of this species to be a generalist, surviving in many different environments. As strains specialize towards life within specific environments, the bacteria no longer require all their cytochrome oxidases and the genes that are no longer required can accrue mutations.

While the role of cytochrome oxidases has been investigated in other pathogens, including *V. cholerae, M. tuberculosis,* and *S. aureus*, the majority of species studied have been facultative anaerobes [9,18–20]. Additionally, most either infect regions like the gut that are expected to have low levels of oxygen, or, like *M. tuberculosis,* have evolved to be able to survive and grow as intracellular pathogens. In these pathogens, individual high-affinity cytochrome oxidases are critical for normal infection. In the respiratory pathogen and obligate aerobe *B. bronchiseptica,* this is not the case. Not only is no individual cytochrome oxidase required for infection, but a predicted low-affinity cytochrome oxidase is sufficient in mice for both the establishment of infection and persistence once bacteria are introduced. Therefore, targeting *bd-*type cytochrome oxidases, which is being explored as a treatment for *M. tuberculosis* among others, would not be effective in *B. bronchiseptica* [51]. This result emphasizes the need to study not only virulence factors but also the basic bacterial physiology to effectively combat bacterial pathogens. By expanding our understanding of the cytochrome oxidases of other pneumonia-causing bacteria, like *Neisseria meningitidis* and *Haemophilus influenzae*, we may find that low-affinity cytochrome oxidases are better drug targets to clear infections within the respiratory tract.

## MATERIALS AND METHOODS

### Bacterial strains, plasmids, and growth conditions

*B. bronchiseptica* strains were grown on Bordet-Gengou (BG) agar plates supplemented with 6% defibrinated sheep blood (Hemostat, catalog no. DSB1) or in Stainer-Scholte (SS) broth supplemented with SS supplement at 37°C ([67], updated in [68]). One strain (*cta1*^+^) had growth defects on BG agar plates and was therefore plated on SS agar plates when possible. As needed, media were supplemented with streptomycin (Sm, 20 μg/mL), gentamicin (Gm, 30 μg/mL), or sucrose (15% w/v). *E. coli* strains were grown in Lysogeny Broth (LB) or on LB agar plates supplemented with diaminopimelic acid (DAP, 300 μg/mL), ampicillin (Ap, 100 μg/mL), and gentamicin (Gm, 30 μg/mL) as needed. All cultures were started from individual colonies from a clonal population.

### Construction of plasmids and strains

The strains and plasmids used in this study are listed in **Table 3** and **Table 4**, respectively, which are based on mutations of the cytochrome oxidase-encoding loci as outlined in **Table 2**. In-frame deletions were generated using allelic exchange via derivatives of the pEG7s vector. We used the DH5α *E. coli* strain for plasmid construction and propagation. We used the Rho3 *E. coli* strain for transforming *B. bronchiseptica*. Any mutations made within *B. bronchiseptica* strains were confirmed by PCR and/or whole-genome sequencing. The order in which deletions were generated in strains with deletions in multiple loci is described in **S2 Figure**.

**Table 2.** Mutations generated in this study.

| Deletion number | Deletion Name | Description |
| --- | --- | --- |
| 1 | $\Delta ccoNOQP$ | Deletion of 3188bp: from nucleotide 7 of *ccoN* (BB3329) to nucleotide 2084 of *ccoP* (BB3326) |
| 2 | $\Delta cydAB1$ | Deletion of 2772bp: from nucleotide 9 of *cydA1* (BB4498) to nucleotide 1147 of *cydB1* (BB4497) |
| 3 | $\Delta cydAB3$ | Deletion of 2373bp: from 41 nucleotides upstream of *cydA3* (BB4012) to nucleotide 994 of *cydB3* (BB4011) |
| 4 | $\Delta cioAB$ | Deletion of 2380bp: from nucleotide 9 of *cioA* (BB1238) to nucleotide 1000 of *cioB* (BB1239) |
| 5 | $\Delta cyoABCD1$ | Deletion of 3842bp: from nucleotide 11 of *cyoA1* (BB1283) to nucleotide 339 of *cyoD1* (BB1286) |
| 6 | $\Delta cyoABCD2$ | Deletion of 3641bp: from nucleotide 6 of *cyoA2* (BB1310) to nucleotide 632 of *cyoC2* (BB1308); *cyoD2* is left intact |
| 7 | $\Delta ctaCDFE1$ | Deletion of 4165bp: from nucleotide 9 of *ctaC1* (BB4831) to nucleotide 493 of *ctaE1* (BB4827) |
| 8 | $\Delta ctaD2$ | Deletion of 1578bp: from nucleotide 10 to nucleotide 1587 of *ctaD2* (BB4674) |

**Table 3.** Plasmids used in this study.

| Plasmid name | Description | Source |
| --- | --- | --- |
| pEG7s | Suicide allelic exchange plasmid for *B. bronchiseptica* used to generate in-frame deletions; confers sucrose sensitivity through *sacB*. Gm<sup>R</sup> Ap<sup>R</sup> | [49] |
| pEG7s-$\Delta ccoNOQP$ | pEG7s containing ~500bp upstream and downstream of mutation 1 | This study |
| pMAB19 | pEG7s containing ~500bp upstream and downstream of mutation 2 | This study |
| pEG7s-$\Delta cydAB3$ | pEG7s containing ~500bp upstream and downstream of mutation 3 | This study |
| pEG7s-$\Delta cioAB$ | pEG7s containing ~500bp upstream and downstream of mutation 4 | This study |
| pEG7s-$\Delta cyo1$ | pEG7s containing ~500bp upstream and downstream of mutation 5 | This study |
| pEG7s-$\Delta cyo2$ | pEG7s containing ~500bp upstream and downstream of mutation 6 | This study |
| PEG7s-$\Delta cta1$ | pEG7s containing ~500bp upstream and downstream of mutation 7 | This study |
| pEG7s-$\Delta ctaD2$ | pEG7s containing ~500bp upstream and downstream of mutation 8 | This study |

**Table 4.** Strains used in this study.

| Strain | Description | Source |
| --- | --- | --- |
| *B. bronchiseptica* |  |  |
| RB50 (WT) | Wild-type *B. bronchiseptica* strain isolated from nares of a naturally infected rabbit; Sm<sup>R</sup> | [55] |
| $\Delta cco$ | RB50 containing mutation 1; Sm<sup>R</sup> | This study |
| $\Delta cyd1$ | RB50 containing mutation 2; Sm<sup>R</sup> | This study |
| $\Delta cyd3$ | RB50 containing mutation 3; Sm<sup>R</sup> | This study |
| $\Delta cio$ | RB50 containing mutation 4; Sm<sup>R</sup> | This study |
| $\Delta cyo1$ | RB50 containing mutation 5; Sm<sup>R</sup> | This study |
| $\Delta cyo2$ | RB50 containing mutation 6; Sm<sup>R</sup> | This study |
| $\Delta cta1$ | RB50 containing mutation 7; Sm<sup>R</sup> | This study |
| $\Delta ctaD2$ | RB50 containing mutation 8; Sm<sup>R</sup> | This study |
| *Bp*-conserved | RB50 containing mutations 1, 3, 4, 6, and 8; Sm<sup>R</sup> | This study |
| *cyo1*<sup>+</sup> *cta1*<sup>+</sup> | RB50 containing mutations 1-4, 6, and 8; Sm<sup>R</sup> | This study |
| *cyd1*<sup>+</sup> *cta1*<sup>+</sup> | RB50 containing mutations 1, 3-6, and 8; Sm<sup>R</sup> | This study |
| *cyd1<sup>+</sup> cyo1<sup>+</sup>* | RB50 containing mutations 1, 3, 4, and 6-8; Sm<sup>R</sup> | This study |
| *cyd1<sup>+</sup>* | RB50 containing mutations 1 and 3-8; Sm<sup>R</sup> | This study |
| *cyo1<sup>+</sup>* | RB50 containing mutations 1-4 and 6-8; Sm<sup>R</sup> | This study |
| *cta1<sup>+</sup>* | RB50 containing mutations 1-6 and 8; Sm<sup>R</sup> | This study |
| *E. coli* |  |  |
| DH5α | Molecular cloning strain derived from K-12 | Gibco-BRL |
| Rho3 | Conjugation strain; Δasd ΔaphA, DAP auxotroph | [69] |

### TTC reduction assay

Cultures of *B. bronchiseptica* were grown overnight at 37°C in SS media supplemented with SS supplement and Sm. Each culture was normalized to 1 OD_600_/mL. Five microliters were spotted onto tryptic soy agar (TSA) plates (Millipore 22091) containing 0.001% (w/v) TTC (2,3,5-triphenyl-tetrazolium chloride) (Sigma-Aldrich T8877) and incubated in the dark for 24 hours at 37°C. Plates were photographed to capture the irreversible color shift from colorless to red that occurs following TTC reduction.

### Liquid culture growth assays

Cultures of *B. bronchiseptica* were grown for 18 hours at 37°C in SS media supplemented with SS supplement and Sm. Each culture was normalized to 0.1 OD_600_/mL and grown on an orbital shaker (VWR Catalog # 89032-096) at 225 rpm at 37°C. Samples were collected at 0, 3, 6, 9, 12, 18, 24, 36, and 48 hours after the start of the experiments. We measured the OD_600_ of these samples and determined the CFU/mL by serially diluting the samples and plating on BG blood agar plates. The number of colonies were enumerated after at least 48 hours growth at 37°C, with *cta1^+^* requiring at least 72 hours growth at 37°C.

### Growth under different atmospheric conditions

Cultures for outgrowth were prepared in the same manner as for growth curves. Samples were then grown on an orbital shaker at 225 rpm at 37 °C within a trigas incubator connected to N_2_ and CO_2_ gas sources (HERACELL VIOS 160i; Fisher Scientific 13998258). To reduce O_2_ levels, pure N_2_ was introduced. We took endpoint samples to ensure that the experimental conditions remained constant throughout and were not disrupted by the reintroduction of atmospheric oxygen at each sampling point. Samples were collected at 24 hours when grown in 5% CO_2_, 48 hours when grown in 5% O_2_ and in 5% O_2_ 5% CO_2_, and 72 hours when grown in 2% O_2_. We measured the OD_600_ of these samples and determined the CFU/mL by serially diluting the samples and plating on BG blood agar plates. The number of colonies were enumerated after at least 48 hours growth at 37°C, with *cta1^+^* requiring at least 72 hours growth at 37°C.

### Bacterial infection of the mouse respiratory tract using the persistence model

Six-week-old female BALB/c mice from Charles River Laboratories (catalog no. BALB/cAnNCrl) were inoculated intranasally with 7.5 × 10^4^ CFU *B. bronchiseptica* in 50 μL of Dulbecco PBS (DPBS). Samples were collected at three hours, one day, three days, seven days, and fourteen days post-infection. At each indicated time point, right lung lobes, trachea, and nasal cavity tissue were harvested from each mouse and the tissues were homogenized in DPBS using a mini-beadbeater with 0.1 mm zirconia beads (Biospec catalog no. 11079110zx). The number of CFU was determined by plating dilutions of tissue homogenates on BG Sm blood agar and enumerating the number of colonies per tissue after at least 48 hours growth at 37°C, with *cta1^+^*requiring at least 72 hours growth at 37°C.

### Bacterial colonization of the mouse respiratory tract

Six-week-old female BALB/c mice from Charles River Laboratories (catalog no. BALB/cAnNCrl) were inoculated intranasally with 100 CFU *B. bronchiseptica* in 4 μL of Dulbecco PBS (DPBS), divided equally between the two nares. Samples were collected at three days and seven days post-infection. At each indicated time point, nasal cavity tissue was harvested and homogenized in DPBS using a mini-beadbeater with 0.1 mm zirconia beads (Biospec catalog no. 11079110zx). The number of CFU was determined by plating dilutions of tissue homogenates on BG Sm blood agar and enumerating the number of colonies per tissue after at least 48 hours growth at 37°C, with *cta1^+^*requiring at least 72 hours growth at 37°C.

### RNA isolation and sequencing

Cultures of *B. bronchiseptica* were grown overnight at 37°C in SS media supplemented with SS supplement and Sm. Each culture was normalized to 1 OD_600_/mL and grown on an orbital shaker at 225 rpm at 37 °C for 4 hours under different atmospheric conditions as above. 3 biological replicates were generated for each condition. Bacteria were pelleted, resuspended in RNAlater (Invitrogen AM7020), then stored at −80°C until extraction. RNA was isolated using Trizol/chloroform extraction, followed by centrifugation in Phasemaker Tubes (Invitrogen 12183555). The aqueous phase was precipitated in isopropanol, washed in ethanol, and solubilized in RNase-free water. Samples were then cleaned using RNAeasy column purification (Quiagen 74104). RNA sequencing was performed by SeqCenter (Pittsburgh, PA; 12M Paired End Reads rRNA depletion sequencing package). Library preparation was performed using Illumina’s Stranded Total RNA Prep Ligation with Ribo-Zero Plus kit and 10bp unique dual indices. Sequencing was performed using a NovaSeq X Plus.

### RNA analysis

RNA analysis was performed by SeqCenter (Pittsburgh, PA; intermediate RNA analysis package). Read mapping was performed using HISAT2 [70] and read quantification was performed using featureCounts [71]. Read counts were loaded into R, then normalized using edgeR’s Trimmed Mean of M values (TMM) algorithm [72]. These values were then converted to counts per million. Differential expression analysis between our different conditions (5% O_2_ vs ambient air, 2% O_2_ vs ambient air, and 5% CO_2_ vs ambient air) was performed using edgeR’s glmQLFTest.

### Ethics statement

All animal studies followed the guidelines in the *Guide for the Care and Use of Laboratory Animals* of the National Institute of Health. Our protocols were approved by the University of North Carolina Institutional Animal Care and Use Committee (Protocol ID: 22-140). All animals were anesthetized for inoculations, monitored daily, and euthanized properly and humanely. All efforts were made to minimize suffering.

## ACKNOWLEDGEMENTS

Thank you to the members of the Cotter lab for their support and the discussions which aided our investigation.

## SUPPORTING INFORMATION

**S1 Table. Genomes of classical bordetellae strains compared**

**S2 Table. Percent amino acid homology of cytochrome oxidases of strains of interest compared to RB50**

**S3 Table. Fold change differences in transcript abundance of all genes of RB50 grown either in 5% O_2_ or ambient air.**

**S4 Table. Fold change differences in transcript abundance of all genes of RB50 grown either in 2% O_2_ or ambient air.**

**S5 Table. Fold change differences in transcript abundance of all genes of RB50 grown either in 5% CO_2_ or ambient air.**

**S1 Figure. Phylogenetic analysis of classical *Bordetella* strains**. This tree was generated by comparing the concatenated multiple sequence alignments in the *cgMLST_genus* scheme in the BIGSdb-Pasteur genomic platform for *Bordetella* [38]. The tree was rooted on the branch leading to *B. petrii*. Leaves are labeled with the strain name and Institut Pasteur Bordetella cgMLST id. More information can be seen in S1 Table. Tree created using Interactive Tree of Life (iTOL) v5 [73].

**S2 Figure. Order of mutations added to generate strains**. A full list of the strains used in this study can be found in Table 4. Numbers represent the in-frame deletion mutation added and also correspond to the mutations listed in Table 2. The names of strains have been added where applicable. Black arrows represent the successful deletion of a cytochrome oxidase-encoding gene locus. Gray crossed lines represent an allelic exchange that resulted only in wild-type revertants; at least 16 clones were screened for each.

**S3 Figure. All generated strains respire**. TTC reduction after 24 hours of growth. When reduced, TTC undergoes an irreversible color change to red. Shown are representative images of 5 biological replicates per strain.

**S4 Figure. The three cytochrome oxidases conserved in *B. pertussis* are sufficient for colonization in mice**. (A) Bacterial burden over time within the nasal cavity (upper), trachea (middle), and right lung (lower) of mice infected with wild-type bacteria (WT, grey circles). Open circles represent tissue samples with no detectable bacteria. n=2 for day 0, n=6 for all other timepoints. (B) Bacterial burden over time within the nasal cavity of mice infected with wild-type bacteria (WT, grey circles) or a strain with only the cytochrome oxidase-encoding gene loci conserved in *B. pertussis* (*Bp-*conserved, maroon hexagon). n=2 for day 0, n=4 for day 3. Each point represents a single mouse. Dashed line represents the limit of detection. Statistical significance was determined using unpaired Student’s t-test; p-values are indicated when p<0.05.

**S5 Figure. No single cytochrome oxidase is required to establish infection.** (A) Growth over time, measured via optical density (left) or CFU/mL (right), for strains with two cytochrome oxidase-encoding gene loci. (B) Bacterial burden over time within the nasal cavity of mice infected with a strain with only the cytochrome oxidase-encoding gene loci conserved in *B. pertussis* (*Bp*-conserved, maroon hexagon), a strain with only *ctaCDFGE1* and *cyoABCD1* (*cta1^+.^ cyo1^+^*, orange upright triangle), a strain with only *cydAB1* and *cyoABCD1* (*cyd1^+^ cyo1^+^*, blue diamond), or a strain with only *cydAB1* and *ctaCDFGE1* (*cyd1^+^ cta1^+^*, green upside-down triangle). Samples from 5 mice were collected at each timepoint for each strain. However, due to the natural microbiota of the nasal cavity, *B. bronchiseptica* could not always be innumerated due to contamination. Therefore, n=4 for *cta1^+^* day 3. Each point represents a single mouse. Dashed line represents the limit of detection. Statistical significance was determined using unpaired Student’s t-test; p-values are indicated when p < 0.05.

## Notes

### Competing Interest Statement

The authors have declared no competing interest.

### Summary of Updates

Supplemental Figures and Tables for the manuscript by McKay et al. have been added.

