## Supplemental Figures for "Cytochrome oxidase requirements in *Bordetella* reveal insights into evolution towards life in the mammalian respiratory tract"

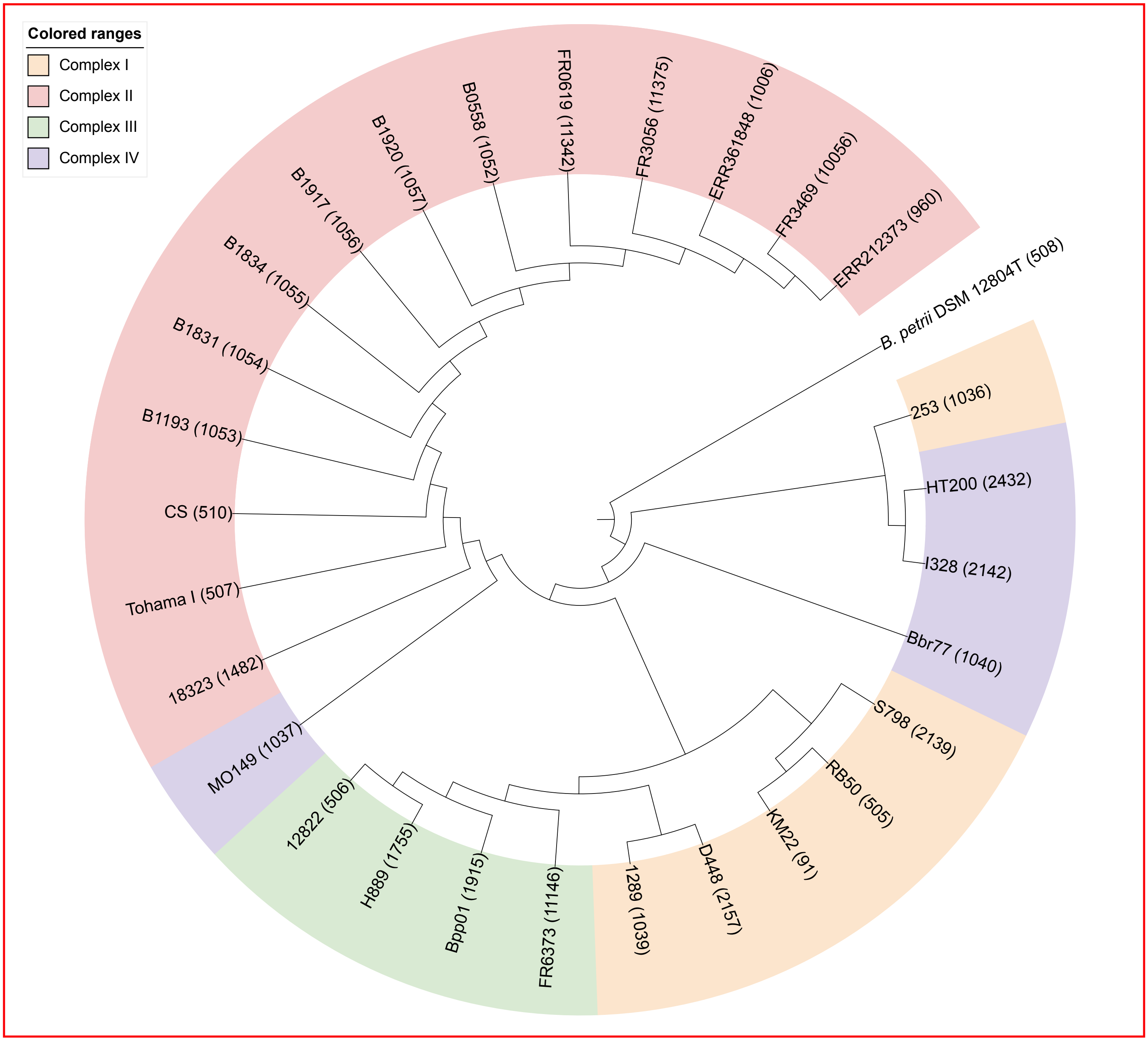


**S1 Figure. Phylogenetic analysis of classical Bordetella strains.** This tree was generated by comparing the concatenated multiple sequence alignments in the cgMLST_genus scheme in the BIGSdb-Pasteur genomic platform for *Bordetella* [38]. The tree was rooted on the branch leading to *B. petrii*. Leaves are labeled with the strain name and Institut Pasteur *Bordetella* cgMLST id. More information can be seen in S1 Table. Tree created using Interactive Tree of Life (iTOL) v5 [74].

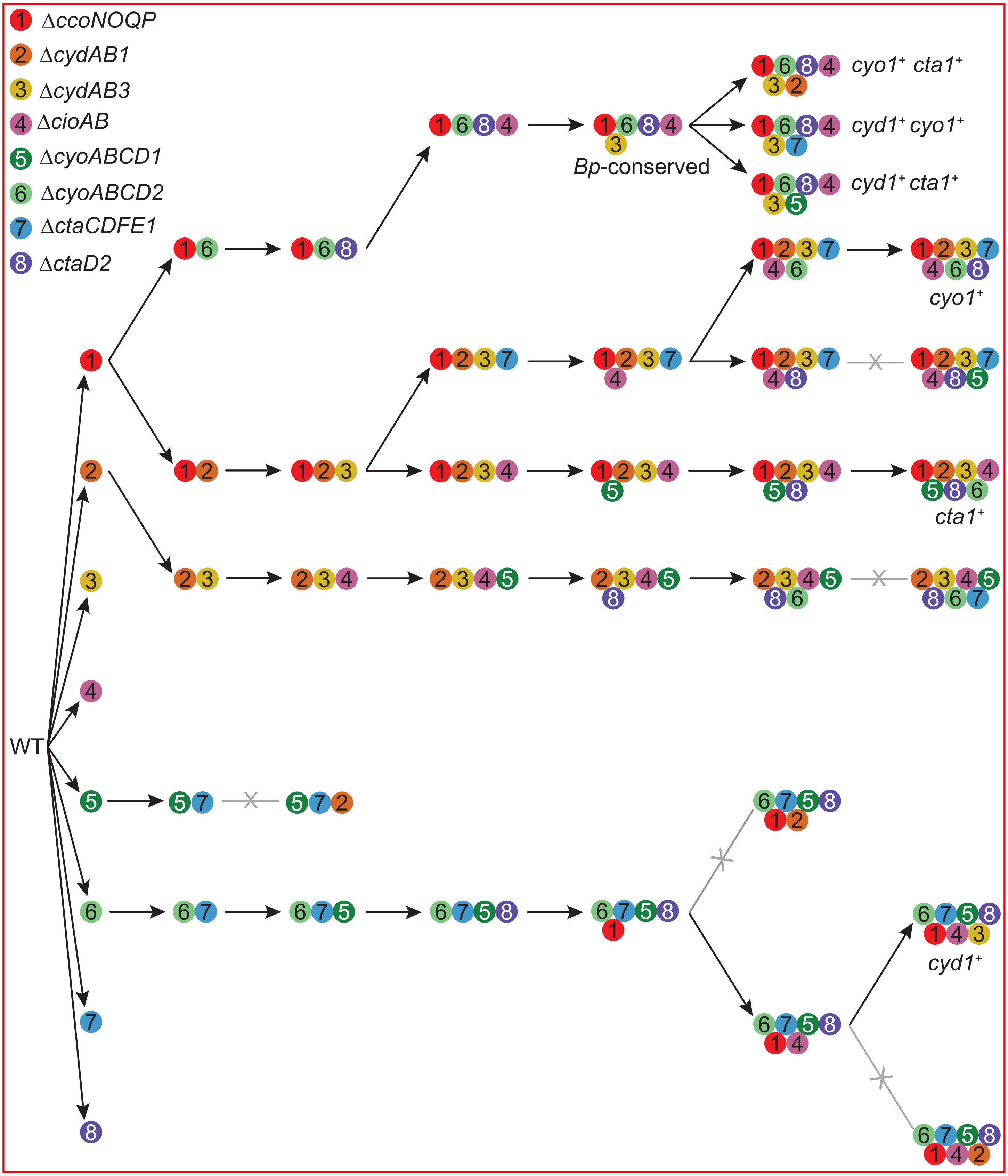


**S2 Figure. Order of mutations added to generate strains.** A full list of the strains used in this study can be found in Table 4. Numbers represent the in-frame deletion mutation added and also correspond to the mutations listed in Table 2. The names of strains have been added where applicable. Black arrows represent the successful deletion of a cytochrome oxidase-encoding gene locus. Gray crossed lines represent an allelic exchange that resulted only in wild-type revertants; at least 16 clones were screened for each.

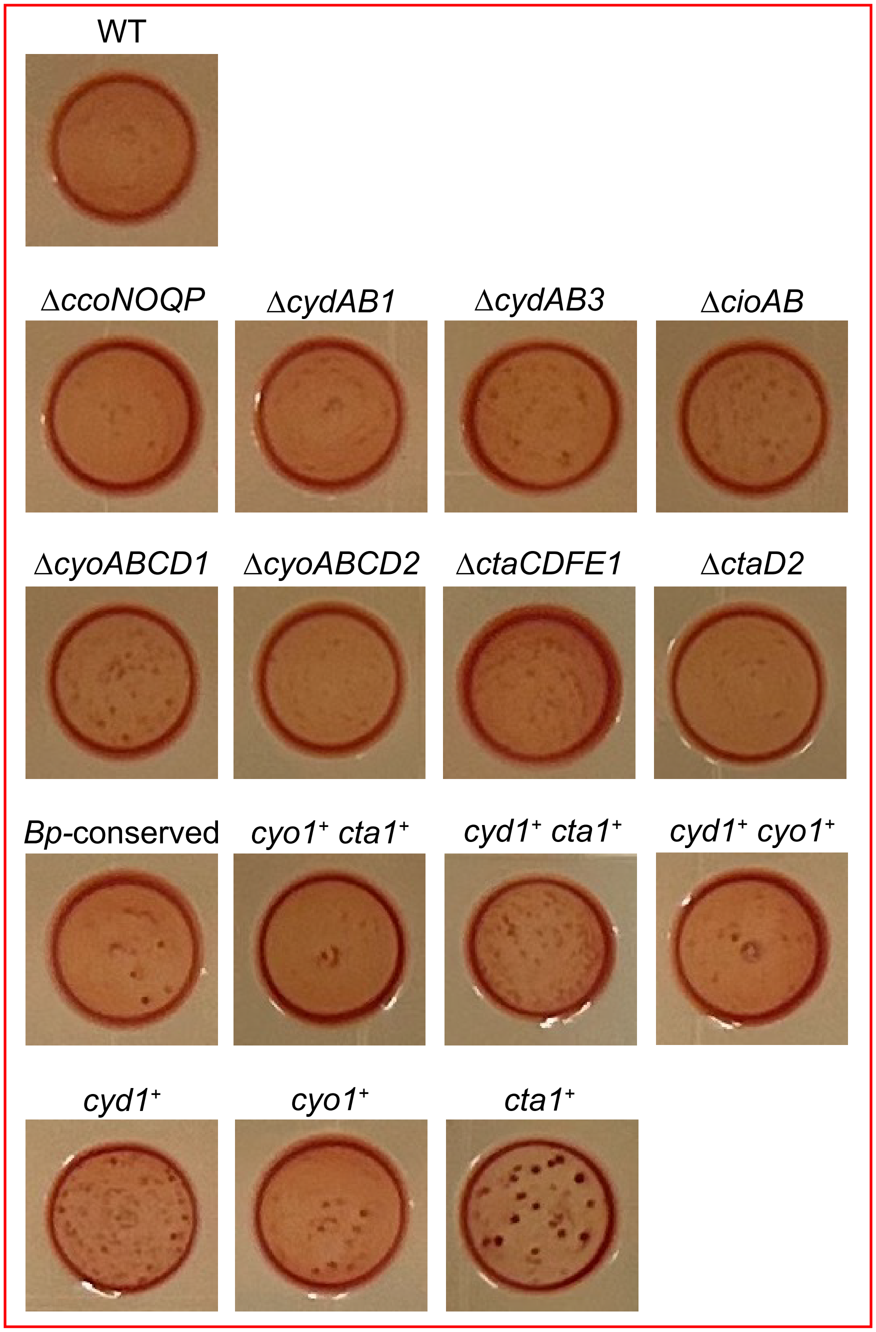


**S3 Figure. All generated strains respire.** TTC reduction after 24 hours of growth. When reduced, TTC undergoes an irreversible color change to red. Shown are representative images of 5 biological replicates per strain.

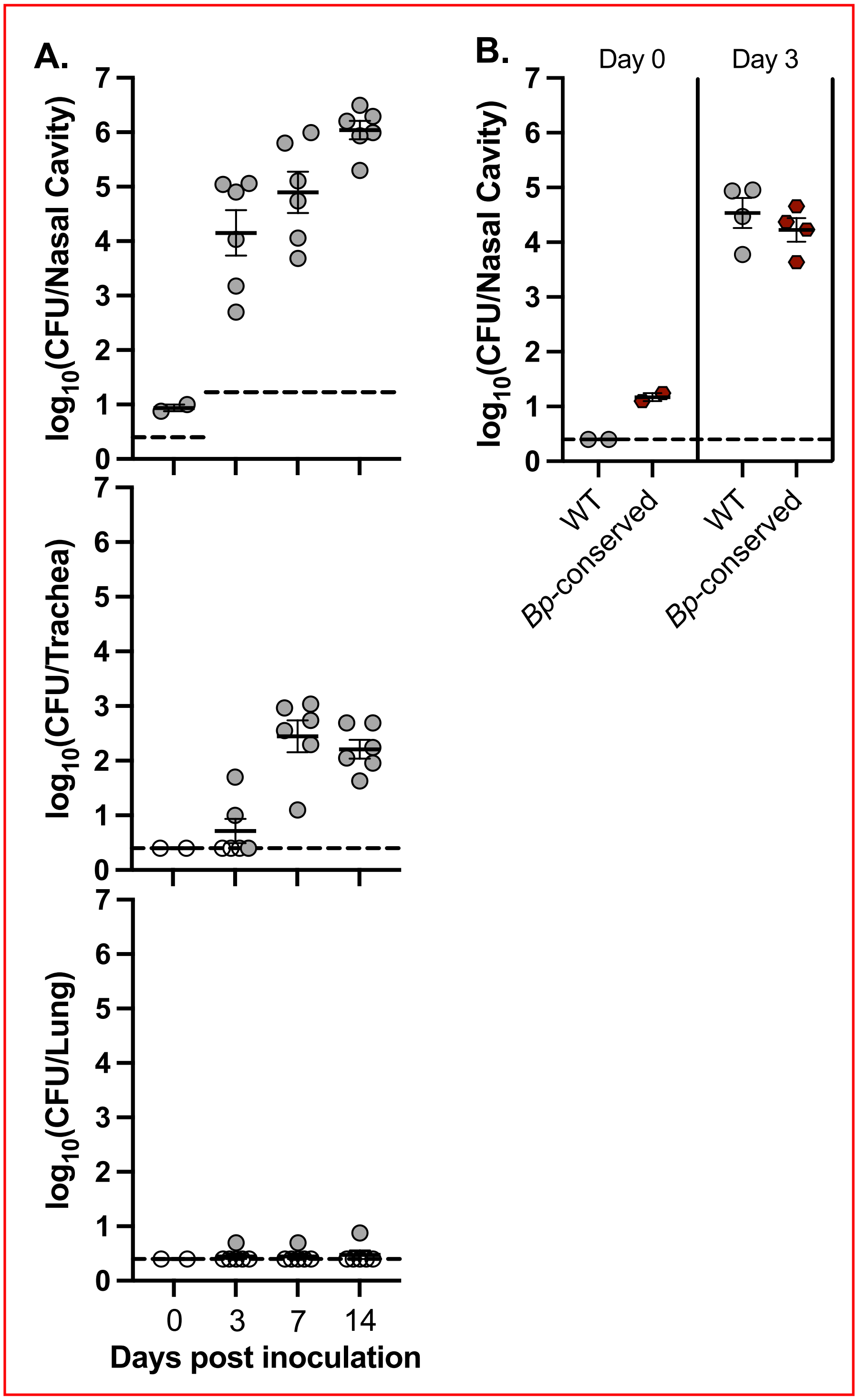


**S4 Figure. The three cytochrome oxidases conserved in *B. pertussis* are sufficient for colonization in mice.** (A) Bacterial burden over time within the nasal cavity (upper), trachea (middle), and right lung (lower) of mice infected with wild-type bacteria (WT, grey circles). Open circles represent tissue samples with no detectable bacteria. n=2 for day 0, n=6 for all other timepoints. (B) Bacterial burden over time within the nasal cavity of mice infected with wild-type bacteria (WT, grey circles) or a strain with only the cytochrome oxidase-encoding gene loci conserved in *B. pertussis* (*Bp*-conserved, maroon hexagon). n=2 for day 0, n=4 for day 3. Each point represents a sing mouse. Dashed line represents the limit of detection. Statistical significance was determined using unpaired Student's t-test; p-values are indicated when $p<0.05$.

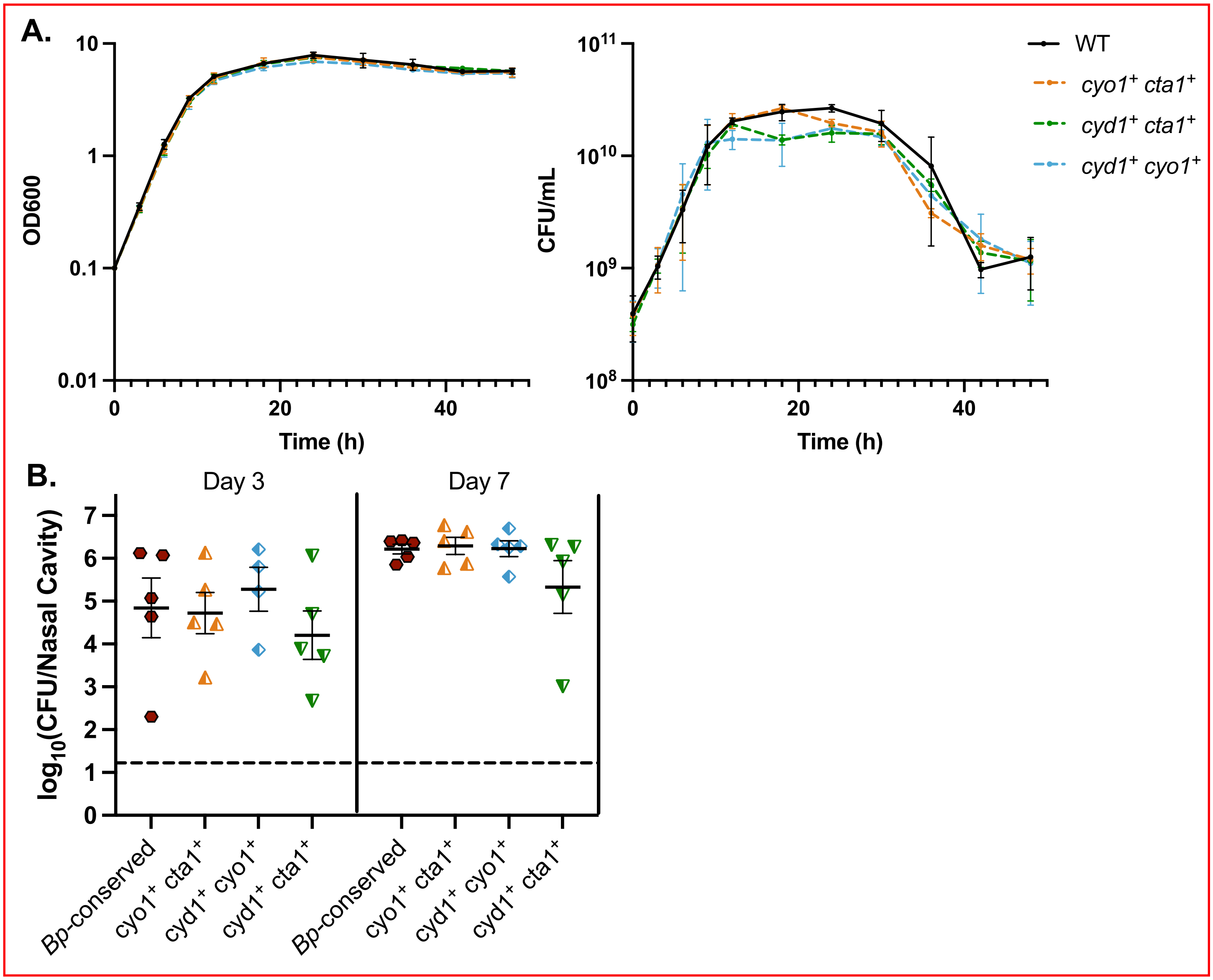


**S5 Figure. No single cytochrome oxidase is required to establish infection.** (A) Growth over time, measured via optical density (left) or CFU/mL (right), for strains with two cytochrome oxidase-encoding gene loci. (B) Bacterial burden over time within the nasal cavity of mice infected with a strain with only the cytochrome oxidase-encoding gene loci conserved in *B. pertussis* (*Bp*-conserved, maroon hexagon), a strain with only *ctaCDFGE1* and *cyoABCD1* (*cta1*$^+$ *cyo1*$^+$, orange upright triangle), a strain with only *cydAB1* and *cyoABCD1* (*cyd1*$^+$ *cyo1*$^+$, blue diamond), or a strain with only *cydAB1* and *ctaCDFGE1* (*cyd1*$^+$ *cta1*$^+$, green upside-down triangle). Samples from 5 mice were collected at each timepoint for each strain. However, due to the natural microbiota of the nasal cavity, *B. bronchiseptica* could not always be innumerated due to contamination. Therefore, n=4 for cta1+ day 3. Each point represents a single mouse. Dashed line represents the limit of detection. Statistical significance was determined using unpaired Student's t-test; p-values are indicated when $p < 0.05$.
